# Scalable saturation mutagenesis reveals gene regulatory architecture and rare variant effects

**DOI:** 10.64898/2026.09.30.755794

**Authors:** Han Yuan, Xingfan Huang, Benjamin Auerbach, Johannes Linder, Divyanshi Srivastava, David R. Kelley

**Affiliations:** Calico Life Sciences LLC, South San Francisco, CA 94080, USA; Institute of Computational Biology, Helmholtz Munich, Neuherberg, Germany; School of Medicine and Health, Technical University of Munich, Munich, Germany

## Abstract

Long-context sequence-to-function models enable nucleotide-resolution prediction of regulatory variant effects, motivating comprehensive mutational interrogation across broad genomic contexts. Yet conventional in silico saturation mutagenesis (ISM) scores mutations one at a time, requiring millions of model evaluations for a single gene and billions to trillions at genome scales. Here, we introduce Multi-ISM, a scalable framework that reformulates ISM as a sparse recovery problem. Multi-ISM generates mutational maps using 45-fold fewer model evaluations than exhaustive single-variant ISM and matches or exceeds its accuracy on variant-effect benchmarks. Multi-ISM is architecture-agnostic and transfers across large-scale sequence-to-function models.

We applied Multi-ISM to 5,000 protein-coding genes, including 3,317 OMIM disease genes, generating base-pair-resolution, tissue-resolved attribution maps across 500-kb windows. These maps supported enhancer–gene prioritization and identification of cell-type-specific regulatory elements. Gene-level summaries of the maps captured regulatory complexity and showed that more constrained genes had smaller predicted mutational effects. Aggregating Multi-ISM predictions into gene-level rare-variant burdens improved personalized expression prediction over a common-variant elastic net, with the largest gains at expression outliers. Multi-ISM makes nucleotide-resolution interpretation of long-context sequence models a routine computation rather than a dedicated effort, so that new architectures, functional readouts, and cellular contexts can be mapped as they appear.

## 2 Introduction

Understanding how genetic variation alters gene regulation is a central challenge in human genetics. Most disease-associated variants lie outside protein-coding regions, where their effects depend on the regulatory elements they perturb, the genes they influence, and the cellular contexts in which the regulation occurs. Sequence-to-function models such as Borzoi and AlphaGenome learn these dependencies directly from genomic sequence and predict molecular phenotypes across many tissues and cell types [1, 2]. These models therefore offer a potential route to systematically estimate regulatory variant effects, interpret the regulatory architecture of individual genes, and prioritize disease-associated variation.

Realizing this potential requires moving beyond scoring predefined sets of observed variants toward comprehensive mutational maps, which estimate the effect of every possible single-nucleotide substitution across the full cis-regulatory context of each gene and across all predicted tissues and cell types. Such maps identify cis-regulatory elements, reveal tissue-specific regulatory architecture, and span the full allele-frequency spectrum, including rare and previously unobserved variants that are difficult to characterize from human cohorts alone. A gene-centered formulation is useful here, because regulatory elements act over hundreds of kilobases and the same nucleotide change can affect different genes differently across cellular contexts.

In silico saturation mutagenesis (ISM) provides the most direct approach for constructing such maps, mutating each nucleotide one at a time and measuring the resulting change in model outputs to yield variant-effect estimates across all predicted cell types. However, conventional ISM scores one substitution per forward pass. Across a 500-kb cis-regulatory window, exhaustive ISM therefore requires roughly 3 million forward passes, given three substitutions per position and reverse-complement augmentation. Applying this procedure genome-wide is costly, even before considering multiple model folds or sequence backgrounds. AlphaGenome Atlas showed that precomputed variant-effect maps at genome scale serve variant prioritization and regulatory interpretation, but building one requires substantial computational resource available to few groups [3]. As newly released sequence-to-function models expand to new architectures, biological readouts, and cellular contexts, there is increasing need for efficient, portable methods to generate comprehensive mutational maps.

Gradient-based methods, including input-gated gradients, integrated gradients, DeepLIFT, DeepSHAP, and Taylor-ISM, rely on backward propagation to estimate nucleotide-level attributions [4–8]. Although efficient, they generally measure local model sensitivity rather than the direct effect of a nucleotide substitution, and require a separate backward pass for each output task, which becomes costly for highly multitask sequence-to-function models.

Other methods accelerate ISM itself. FastISM avoids recomputing convolutional activations outside the receptive field affected by a mutation [9], whereas Yuzu compresses the sparse differences between mutant and reference activations before convolution and reconstructs them afterward through sparse recovery [10]. Both closely reproduce conventional ISM but are difficult to extend to long-range attention, complex residual operations, and other unsupported layers. Surrogate approaches such as SQUID and Multiplexer remain limited to short-context models [11, 12]. Orca introduced multiplexed in silico perturbation, but focused on short-sequence disruptions affecting 3D genome structure [13]. Thus, there remains a need for an efficient saturation-mutagenesis framework that scales to broad regulatory contexts while remaining independent of the internal architecture of the underlying model.

Here, we introduce **Multi-ISM**, an architecture-agnostic framework for efficient in silico saturation mutagenesis across broad genomic contexts. Multi-ISM exploits the sparsity of regulatory effects. Although millions of substitutions are possible within a 500-kb window, only a small fraction meaningfully alter the model prediction of a given gene. Thus, Multi-ISM introduces many spatially separated mutations in each forward pass, measures their aggregate effect, and estimates individual variant effects through sparse signal recovery. Efficiency is further improved through an adaptive strategy that iteratively prunes resolved high-effect mutations and refines the remaining unresolved variants. Because Multi-ISM requires only model inputs and outputs, the same framework can be applied across distinct sequence-to-function architectures and readouts.

We show that Multi-ISM recovers informative variant effects with substantially fewer model evaluations than conventional single-variant ISM, preserves experimentally supported regulatory features, and filters variants with little detectable signal. We further demonstrate that Multi-ISM transfers across transformer- and state-space-based sequence-to-function models, extending comprehensive mutational analysis to additional cellular contexts. Applying Multi-ISM to 5,000 genes, we characterize regulatory architectures ranging from compact, shared programs to complex tissue-specific landscapes. After accounting for expression, tissue specificity and gene structure, we find that more constrained genes have smaller predicted mutational effects and more diverse regulatory patterns, consistent with regulatory buffering. By aggregating predicted rare-variant effects into gene-level burdens, Multi-ISM improves personalized gene-expression prediction, with the largest gains at expression outliers. Together, these results establish Multi-ISM as a scalable and portable framework for dissecting gene regulatory architecture and rare regulatory variation.

## 3 Results

### 3.1 Multi-ISM reformulates saturation mutagenesis as a sparse recovery problem

In silico saturation mutagenesis (ISM, or single-ISM) evaluates one mutation per forward pass, so the number of model evaluations scales linearly with sequence length. This becomes expensive for long-context sequence-to-function models, where exhaustive mutagenesis across a 500-kb input window requires millions of model evaluations. However, most mutations across such a window have little or no predicted effect on target gene expression, because regulatory elements are sparsely distributed in the genome. Moreover, current sequence-to-function models do not reliably predict very small effects; sign prediction by both Borzoi and AlphaGenome approaches random for fine-mapped eQTLs with predicted magnitude below 10*^−^*^3^ (**Supp. Fig. S1**). Thus, exhaustively evaluating the effect of every single mutation is highly inefficient.

Multi-ISM exploits this sparsity to reduce the cost of variant effect estimation. Multi-ISM introduces multiple spatially separated mutations into each sequence, measures their aggregate effect on model output, and reconstructs individual variant effects by solving a sparse linear system. Mutations within the same sequence are separated by a minimum distance to preserve approximate additivity. We refer to this implementation as base Multi-ISM (**Methods, Fig. 1A**).

**Figure 1:**
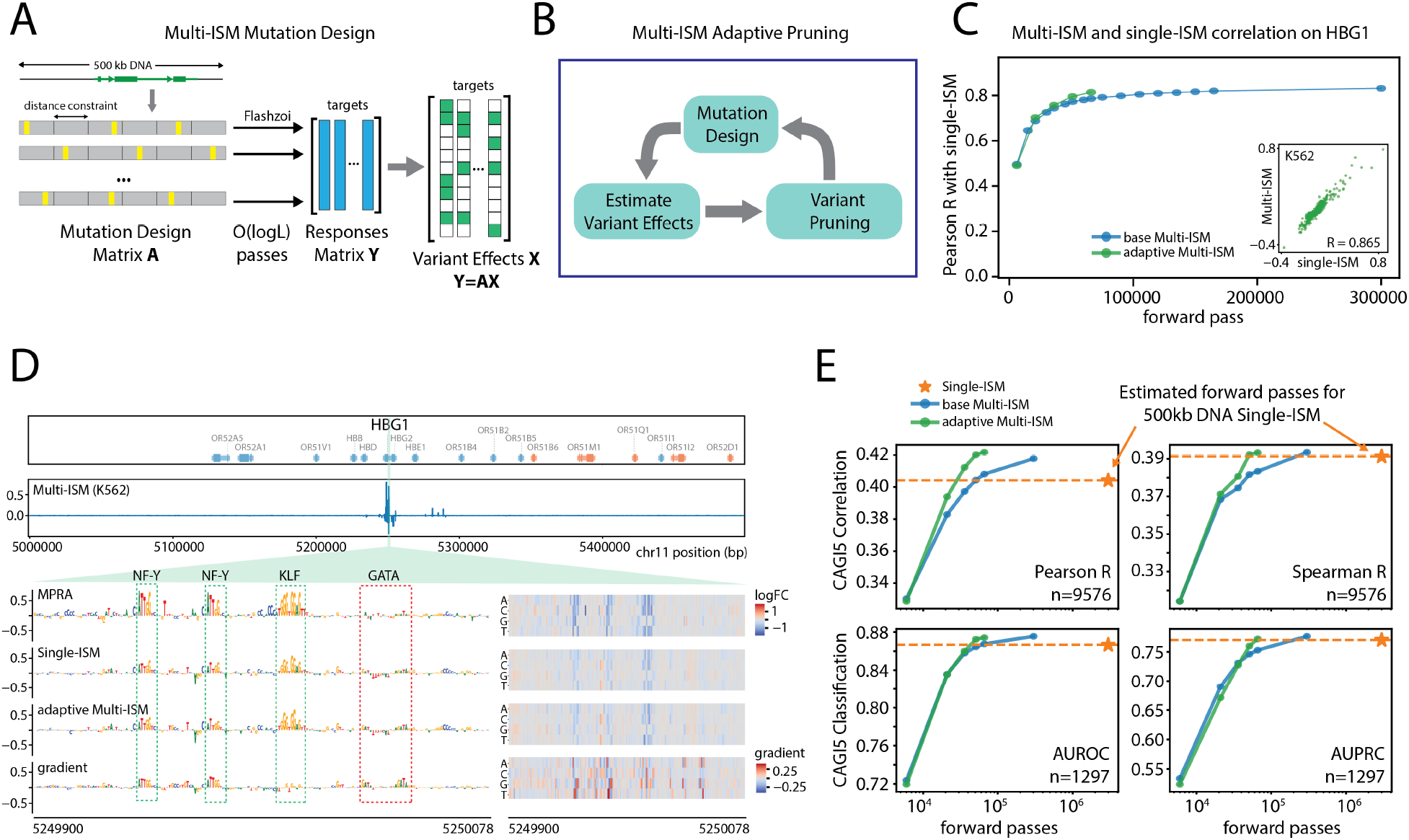
Multi-ISM makes efficient and accurate variant-effect predictions. **A,** Schematic of base Multi-ISM. Multiple spatially separated mutations are introduced into each sequence, and individual variant-effect estimates are recovered from the resulting multi-mutant measurements by sparse regression. **B,** Schematic of adaptive Multi-ISM. Variant effects are iteratively estimated via regression, high-effect variants are re-evaluated and pruned from subsequent designs, and the remaining variants are re-estimated using additional Multi-ISM measurements. **C,** Pearson correlation between Multi-ISM and single-ISM predictions for HBG1 averaged across output tracks, as a function of the number of forward passes. The comparison includes variants within the central 10-kb window with absolute single-ISM effects greater than 0.01. Blue, base Multi-ISM; green, adaptive Multi-ISM. Inset, representative comparison of adaptive Multi-ISM and single-ISM scores averaged across K562 tracks. **D,** Top, adaptive Multi-ISM attribution scores across the 500 kb region surrounding HBG1 in K562. In gene tracks, orange and blue denote plus- and minus-strand genes. Bottom, CAGI5 MPRA measurements, single-ISM, adaptive Multi-ISM and Input *×* Gradient attribution scores near HBG1 promoter. Dashed boxes highlight regulatory motifs. Nucleotide-level substitution effects are shown at right for the indicated region; for gradient track, nucleotide-level gradient scores are shown. **E,** Performance of base Multi-ISM (blue) and adaptive Multi-ISM (green) on the CAGI5 MPRA saturation mutagenesis benchmark as a function of computational budget. Pearson and Spearman correlations quantify agreement with measured variant effects, whereas AUROC and AUPRC quantify classification of over-expression versus under-expression variants as defined previously (**Methods**). Dashed orange lines indicate single-ISM performance, and orange stars indicate the estimated cost of single-ISM across a 500 kb sequence. *N* indicates the number of variants used for regression or classification.

We used Flashzoi, a transfer-learned re-implementation of Borzoi with a more efficient transformer backbone, to ask whether base Multi-ISM could approximate single-ISM variant-effect estimates [1, 14]. Starting with genes in the CAGI5 MPRA saturation mutagenesis dataset, we computed base Multi-ISM across 500-kb windows and compared its estimates with single-ISM (**Methods**). Exhaustive single-ISM across 500 kb requires 3 million forward passes, accounting for three alternative mutations per position and forward/reverse-complement augmentation. We tuned key Multi-ISM hyperparameters (**Methods**), and found that agreement with single-ISM improved as the number of measurements increased, approaching a plateau at most loci by 300,000 forward passes, or 10% of the cost of full-window single-ISM (**Fig. 1C**, **Supp. Fig. S1**). For example, at 300,000 forward passes, agreement with single-ISM for HBG1 reached Pearson’s R=0.831 and Spearman’s R=0.670, averaged across all RNA and CAGE tracks. These results show that sparse recovery can approximate single-ISM effect maps with substantially fewer model evaluations.

We next examined the assumptions and scaling behavior of base Multi-ISM. Under our spatially separated multi-mutant design, adding a global nonlinearity to the additive model did not improve held-out prediction (**Methods, Supp. Fig. S2**). Multi-ISM also remained effective as the mutagenized region expanded. The performance gap between 100-kb and 500-kb windows, where five-fold more variants must be estimated, was largest at 15,000 measurements but nearly closed by 250,000 (**Supp. Fig. S2**). This scaling is expected for sparse recovery, where measurement requirements depend primarily on the number of non-zero effects rather than the total number of variants.

To further improve efficiency, we introduce an adaptive pruning strategy. We observed that a few high-effect variants dominate each multi-mutant measurement and leave large residual errors that obscure weaker effects. Adaptive Multi-ISM therefore iteratively identifies these dominant variants, re-estimates each one individually across randomly mutated sequence backgrounds, and removes them from subsequent mutation designs and regression fits (**Methods, Fig. 1B, Supp. Fig. S1**).

Adaptive Multi-ISM achieved greater correlation with single-ISM than base Multi-ISM across genes at given computing budgets (**Fig. 1C**, **Supp. Fig. S1**). For HBG1, adaptive Multi-ISM reached Pearson’s R=0.814, Spearman’s R=0.601 with single-ISM using only 66,000 forward passes, more concordant than base Multi-ISM at the same budget (Pearson’s R=0.786, Spearman’s R=0.528; **Fig. 1C**).

We next examined whether Multi-ISM preserves biologically interpretable regulatory features. On the HBG1 promoter, where the CAGI5 MPRA data identify variants disrupting known regulatory elements, both single-ISM and adaptive Multi-ISM recovered strong effects at NFY and KLF motifs, consistent with the MPRA measurements (**Fig. 1D**). In contrast, the gradient failed to highlight the KLF site and instead assigned high saliency to a nearby GATA motif not supported by the MPRA data. At this locus, Multi-ISM recovered the motif-level signal detected by single-ISM and supported by the MPRA measurements more faithfully than gradient-based attribution.

Although agreement between adaptive Multi-ISM and single-ISM increased with computational budget, the two methods did not converge to perfect agreement (**Fig. 1C**), indicating differences in their variant-effect estimates. We therefore compared their accuracy against experimentally measured effects across the CAGI5 MPRA dataset. Across both regression and classification tasks, Multi-ISM approaches matched or modestly exceeded single-ISM (**Fig. 1E**). Generating a full 500-kb map required 66,000 forward passes for adaptive Multi-ISM and 300,000 for base Multi-ISM, compared with an estimated 3,000,000 for exhaustive single-ISM; for this analysis, single-ISM benchmark scores were computed only for variants within the assayed elements. For example, despite only moderate agreement with single-ISM for LDLR (Pearson’s R = 0.443), adaptive Multi-ISM better matched experimental measurements across both correlation and classification metrics (Pearson’s R: 0.409 vs. 0.366; Spearman’s R: 0.263 vs. 0.241; AUROC: 0.711 vs. 0.694; AUPRC: 0.479 vs. 0.447, adaptive Multi-ISM vs. single-ISM; **Supp. Fig. S1**). One possible explanation is that Multi-ISM estimates each variant across multiple sequence backgrounds rather than only the reference sequence, reducing dependence on that single background. Because adaptive Multi-ISM matched base Multi-ISM on this benchmark using more than 4-fold fewer forward passes, we use the adaptive implementation in subsequent analyses and refer to it simply as Multi-ISM.

### 3.2 Sparse recovery selects the eQTL variants with predictable effects

To further evaluate Multi-ISM versus single-ISM on variant effect prediction, we used the 1,778 protein-coding genes with Multi-ISM maps that harbor at least one fine-mapped eQTL (PIP *≥* 0.9) in one of the 26 GTEx tissue groups analyzed, yielding 6,712 variant–gene–tissue triplets (**Methods**). Multi-ISM effects were read from the precomputed maps, whereas single-ISM here means scoring each eQTL variant individually, rather than performing mutagenesis over a window. Overall, the two methods are highly concordant (Pearson’s *R* = 0.953; **Fig. 2A**). Multi-ISM assigns non-zero effects to 36.1% of these variants in the corresponding tissue (selected variants), while the remaining 63.9% are zeroed out (filtered variants; **Fig. 2B**).

**Figure 2:**
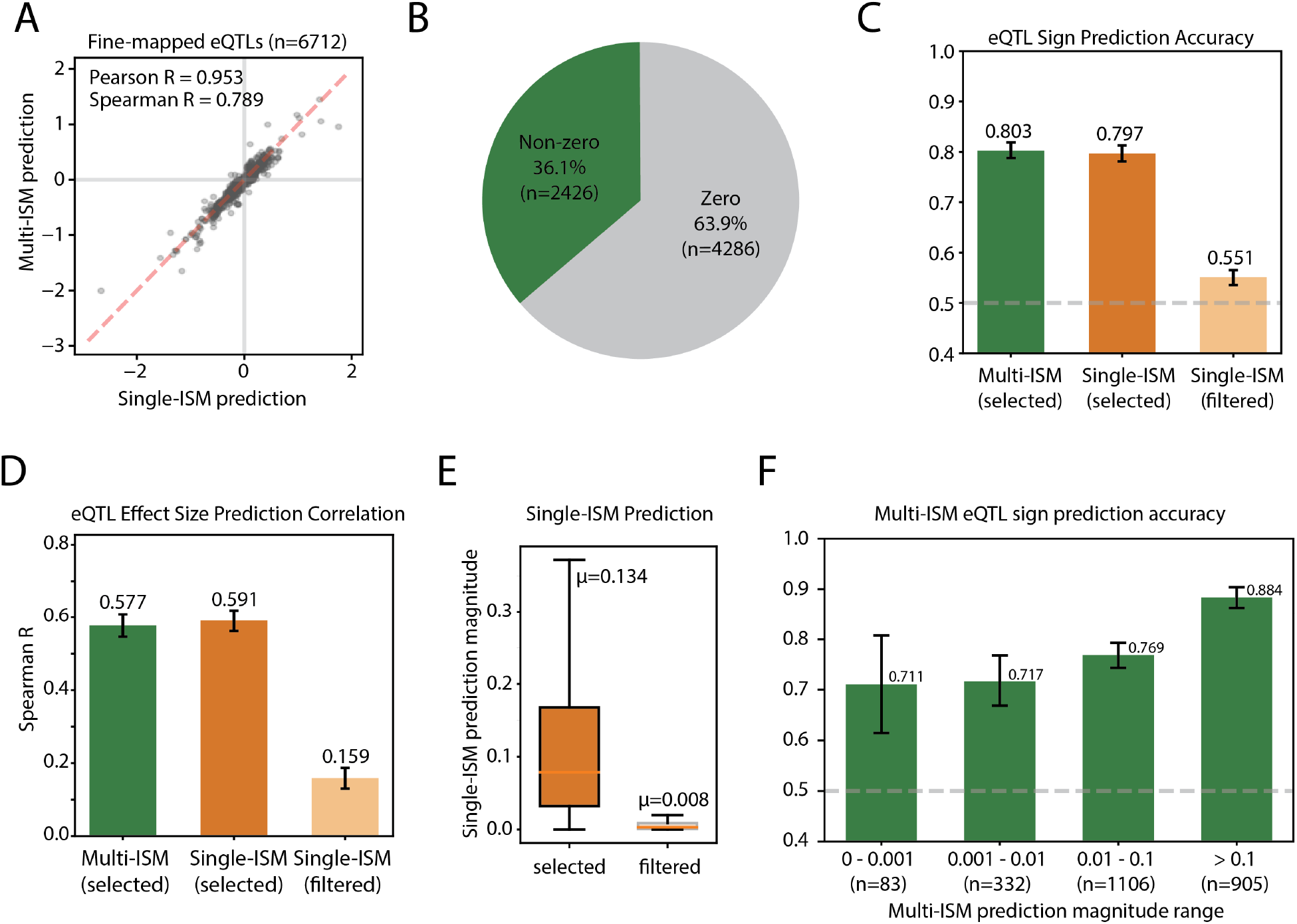
Multi-ISM sparse recovery retains informative regulatory variants. **A,** Correlation of Multi-ISM and single-ISM predictions for 6,712 fine-mapped (PIP*≥*0.9) GTEx v8 eQTL variant–gene–tissue triplets. **B,** Percentage of eQTL variants assigned a non-zero effect by Multi-ISM (selected; 36.1%, n=2,426) or shrunk to zero (filtered; 63.9%, n=4,286). **C,** eQTL sign-prediction accuracy for the 2,426 variants selected by Multi-ISM, scored using either Multi-ISM (green) or single-ISM (orange), and for the 4,286 variants filtered by Multi-ISM, scored using single-ISM (light orange). Error bars indicate 95% confidence intervals estimated from 1,000 bootstrap resamples. **D,** Spearman correlation between predicted and measured eQTL effect sizes for the same groups as in (C). **E,** Distribution of single-ISM predicted magnitude for variants selected or filtered by Multi-ISM. The mean of each distribution is indicated (*µ*=0.134 and 0.008, respectively). **F,** Multi-ISM eQTL sign prediction accuracy stratified by predicted magnitude. Numbers below bars indicate the variants evaluated in each bin. Error bars indicate 95% confidence intervals estimated from 1,000 bootstrap variant resamples.

For selected variants, Multi-ISM and single-ISM predictions were highly consistent (Pearson’s *R* = 0.957; Spearman’s *R* = 0.956; **Supp. Fig. S3**), and the two methods achieved nearly identical performance in eQTL sign prediction (accuracy 0.803 for Multi-ISM vs. 0.797 for single-ISM; **Fig. 2C**) and effect-size prediction (Spearman’s *R* = 0.577 vs. 0.591; **Fig. 2D**). By contrast, single-ISM predicted very small effects for the filtered variants, with a mean absolute log_2_ fold change of 0.008 (**Fig. 2E**), and their single-ISM sign accuracy was only slightly above chance (0.551; **Fig. 2C**). Thus, variants filtered by Multi-ISM are predominantly those for which single-ISM also provides little predictive signal.

As noted above, Borzoi and AlphaGenome variant effect prediction performance (single-ISM) declines sharply at lower predicted magnitudes, approaching random below 10*^−^*^3^ (**Supp. Fig. S1**). Stratifying Multi-ISM eQTL predictions by predicted magnitude, we found that the variants Multi-ISM retains remain informative even at low predicted magnitudes: among the 83 selected variants with predicted magnitude below 10*^−^*^3^, sign accuracy was 0.711 (**Fig. 2F**). Scoring the same set with single-ISM gave a comparable sign accuracy of 0.735, so the retained signal reflects which variants Multi-ISM selects rather than more accurate effect estimates (**Supp. Fig. S3**).

### 3.3 Multi-ISM generalizes across sequence-to-function models and variant-effect readouts

Since Multi-ISM operates only through model inputs and outputs, it does not depend on the internal architecture of the underlying sequence-to-function model. To test this portability, we applied the adaptive Multi-ISM framework to two additional architectures, Borzoi-Prime and Cerberus (**Fig. 3A**). Borzoi-Prime is a transformer-based model with a 524 kb input window and 5,432 output tracks [15]; we applied adaptive Multi-ISM over a 500 kb mutational window to compute variant-effect scores for 2,444 CAGE, RNA-seq, and scRNA-seq expression tracks. Cerberus uses Hydra state-space blocks in place of transformers and has a 786 kb input window with 8,361 output tracks [16]; accordingly, we expanded the mutational window to 700 kb and scored variants across 3,171 CAGE, RNA-seq, and scRNA-seq expression tracks.

**Figure 3:**
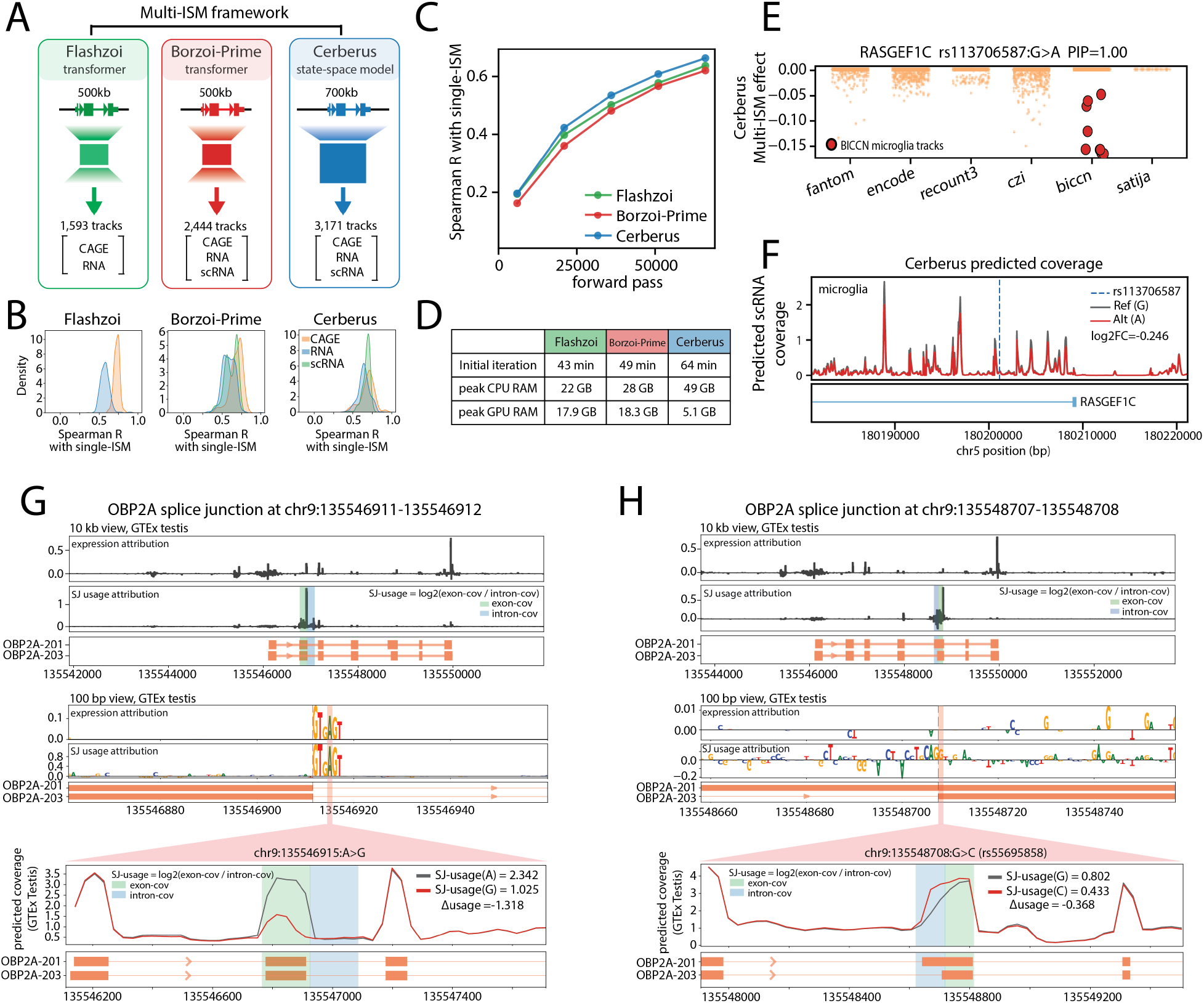
Multi-ISM generalizes across sequence-to-function models and variant-effect readouts. **A,** Multi-ISM was applied to Flashzoi, a transformer model with a 500 kb mutational window and 1,593 CAGE/RNA tracks; Borzoi-Prime, a transformer model with a 500 kb mutational window and 2,444 CAGE/RNA/scRNA tracks; and Cerberus, a state-space model with a 700 kb mutational window and 3,171 CAGE/RNA/scRNA tracks. **B,** Distribution of Spearman correlations between Multi-ISM and single-ISM variant-effect predictions across output tracks for the example gene *GP1BB*. The comparison includes variants within the central 10-kb window with absolute single-ISM effects greater than 0.01. Additional genes are shown in **Supp. Fig. S4**. **C,** Spearman correlation between Multi-ISM and single-ISM predictions for *GP1BB*, averaged across all tracks, as a function of the number of forward passes. Green, Flashzoi; red, Borzoi-Prime; blue, Cerberus. **D,** Computational requirements for Multi-ISM across the three models measured on NVIDIA RTX 4090 GPU. Initial iteration denotes the wall-clock time for the first Multi-ISM iteration comprising 6,000 forward passes. Peak CPU and GPU memory usage are also shown. **E,** Cerberus Multi-ISM effects for the fine-mapped *RASGEF1C* eQTL rs113706587:G*>*A across Cerberus output tracks. BICCN microglia tracks are highlighted in red. **F,** Cerberus-predicted scRNA-seq coverage across *RAS-GEF1C* for the reference (G, gray) and alternative (A, red) alleles at rs113706587, averaged across BICCN microglia tracks (mean single-ISM logFC=*−*0.246). The dashed blue line marks the variant position. **G– H,** Flashzoi Multi-ISM attribution for expression and splice-junction usage in the GTEx testis track at two *OBP2A* splice-site variants: chr9:135,546,915:A*>*G near a splice donor (G) and chr9:135,548,708:G*>*C (rs55695858) at a splice acceptor (H). Expression is the standard readout, defined as the log of summed predicted exonic coverage; splice-junction usage (SJ usage) is defined as the log_2_ ratio of exon-side (green) to intron-side (blue) predicted coverage. Each example shows a 10 kb locus view, a 100 bp nucleotide-resolution view around the highlighted variant (red), and Flashzoi-predicted coverage for the reference and alternate outs. Consistent with this, Borzoi predicted reduced donor usage (Δusage = *−*1.318), increased exon skipping, and decreased transcript abundance (Fig. 3G). By contrast, chr9:135,548,708:G*>*C (rs55695858), a ClinVar variant reported to weaken a splice acceptor (accession: VCV004353954.1), showed strong SJ-usage attribution but little expression attribution. Borzoi predicted reduced usage of the annotated acceptor and a shift toward an upstream acceptor (Δusage = *−*0.368), with minimal change in total transcript abundance (Fig. 3H). We observed a similar pattern for PAS usage at *SRSF11*, where variants altered PAS usage while having little effect on total expression (**Supp. Fig. S5**). Together, these examples show that the same Multi-ISM forward passes used to map expression effects can reveal SJ- and PAS-usage effects at no additional model-evaluation cost. These readouts distinguish transcript-processing effects that alter total abundance from those largely invisible to expression-based attribution.

Using the same hyperparameters and forward-pass budget across architectures, Multi-ISM achieved agreement with single-ISM for Borzoi-Prime and Cerberus comparable to that for Flashzoi. For the example gene *GP1BB*, median Spearman correlations were 0.73 and 0.57 for Flashzoi CAGE and RNA tracks; 0.69, 0.58, and 0.62 for Borzoi-Prime CAGE, RNA, and scRNA tracks; and 0.70, 0.64, and 0.69 for the corresponding Cerberus tracks (**Fig. 3B**). Agreement varied across genes, but for all three architectures it increased with additional forward passes (**Fig. 3C**, **Supp. Fig. S4**). Cerberus agreement remained competitive under the same forward-pass budget despite a 700-kb mutational window, 40% larger than the 500-kb windows used for Flashzoi and Borzoi-Prime. This is consistent with distal regulatory effects becoming increasingly sparse with distance from the target gene, such that expanding the mutational window adds relatively few additional non-zero effects to recover. Computational requirements remained practical across models (**Fig. 3D**), and model-specific hyperparameter tuning may further improve performance.

Applying Multi-ISM across models enables variant-effect mapping in additional cellular contexts. For example, extending Multi-ISM to Cerberus allowed us to score variants across BICCN brain cell-type tracks. For the fine-mapped *RASGEF1C* microglia eQTL rs113706587:G*>*A (PIP=1.0), which colocalizes with Alzheimer’s disease (PP4=1.0) [17], Multi-ISM predicted a strong reduction in microglial expression that was largely absent from other BICCN cell-type tracks and most bulk RNA-seq and CAGE tracks (**Fig. 3E-F**). Together, these results demonstrate that Multi-ISM transfers across long-context sequence-to-function architectures with minimal modification and can extend variant-effect mapping into new models and new cellular contexts.

Beyond model architecture, Multi-ISM is also agnostic to how predicted coverage is summarized into variant-effect readouts. Our primary analyses summarize changes in total predicted exonic coverage. The coverage predictions from each forward pass can also be used to derive splice-junction (SJ) and polyadenylation-site (PAS) usage effects with the same regression framework, without additional model evaluations (**Methods**). As a proof of principle, we computed SJ and PAS usage scores for representative genes (**Fig. 3G–H**; **Supp. Fig. S5**; Methods).

At *OBP2A*, chr9:135,546,915:A*>*G lies near a splice donor and is reported in ClinVar to promote exon skipping (accession: VCV004353946.1). This variant is predicted to perturb both splicing and overall transcript abundance, and so it showed strong Multi-ISM attribution in both read-

### 3.4 Multi-ISM maps prioritize enhancers and resolve their tissue specificity

We next used the Flashzoi Multi-ISM maps generated across thousands of genes to characterize cis-regulatory architecture over long genomic contexts. We first asked whether regulatory elements could be reliably identified from these maps by evaluating whether Multi-ISM attribution scores could prioritize functional enhancer–gene links using a CRISPRi-FlowFISH benchmark. Across 53 genes, we scored enhancer–gene pairs from the attribution maps and compared them with Enformer, Borzoi gradient, and Flashzoi gradient scores by AUROC and AUPRC, stratified by TSS distance. Multi-ISM achieved the highest AUPRC in every distance bin (**Fig. 4A**). However, the 95% confidence intervals overlapped across methods, and AUROC performance varied more across bins (**Supp. Fig. S6**). These results support the use of Multi-ISM attribution scores to prioritize functional enhancer–gene links.

**Figure 4:**
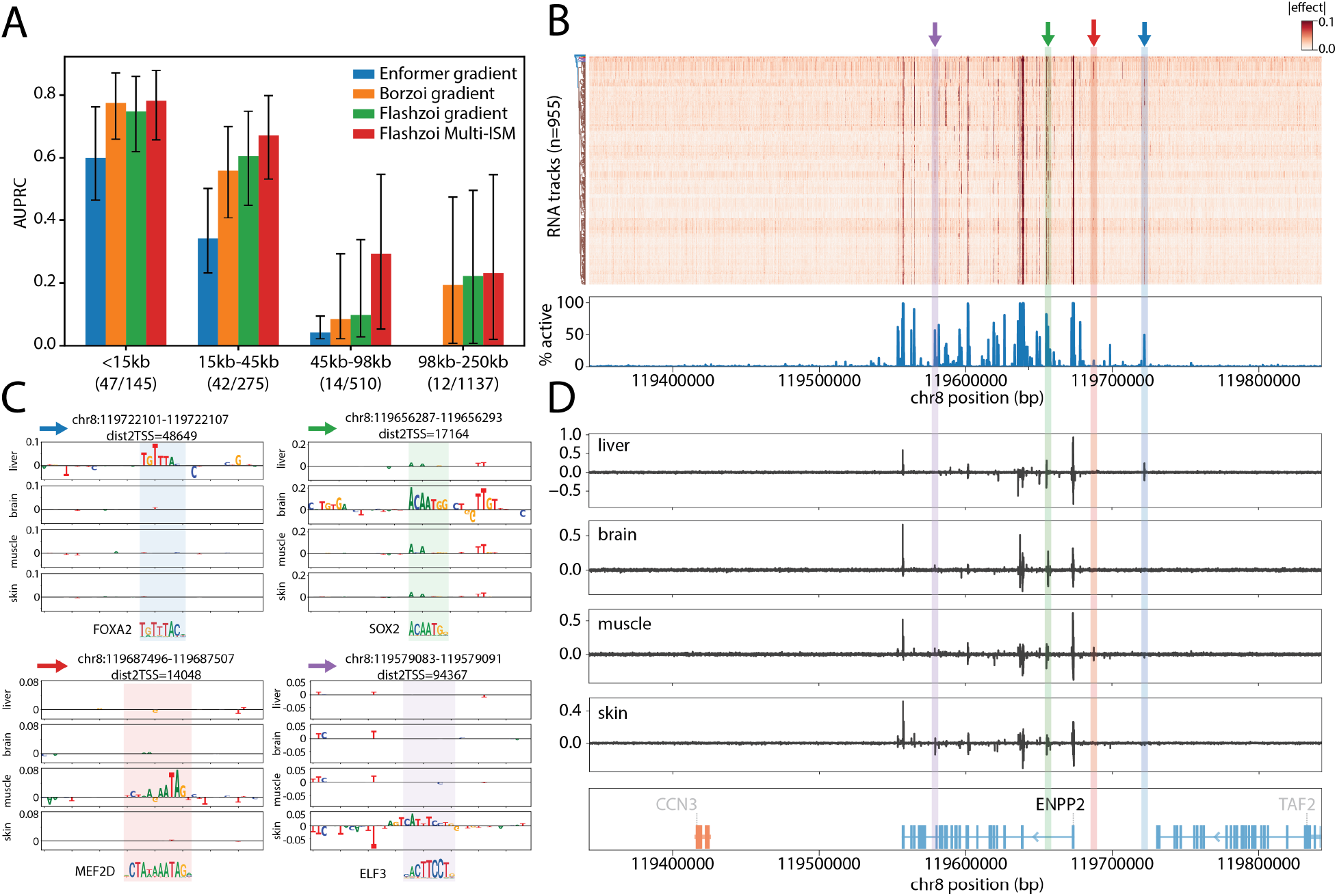
Multi-ISM reveals tissue-specific regulatory architecture. **A,** AUPRC on the K562 CRISPRi-FlowFISH enhancer prioritization benchmark across 53 genes, stratified by enhancer–TSS distance (*<*15 kb, 15–45 kb, 45–98 kb, and 98–250 kb). For Enformer, we used performance reported in the Borzoi study; the 196,608 bp input sequence excludes the most distant bin. For Borzoi gradients, Flashzoi gradients, and Multi-ISM (Flashzoi-based), we report fold-0 performance for a matched comparison. Numbers below each bin indicate positive/total enhancer–gene pairs tested. Error bars indicate 95% confidence intervals estimated from 1,000 bootstrap resamples. **B,** Multi-ISM regulatory map across the 500-kb window surrounding *ENPP2* for 955 RNA tracks. Top, track-by-position attribution heatmap, with each 100-bp bin summarized by its maximum absolute attribution. Tracks are ordered by hierarchical clustering (dendrogram at left). Bottom, percentage of tracks in which each 100-bp bin is active, defined as a maximum absolute attribution *>* 0.05. Colored arrows and shaded regions mark four candidate tissue-selective regulatory elements examined in (C) and (D). **C,** Seqlogo view of base-pair resolution Multi-ISM attribution at the four candidate ENPP2 regulatory elements highlighted in (B), shown for liver (blue), brain (green), muscle (red) and skin (purple). Matched motifs shown below. dist2TSS means distance to most upstream TSS. **D,** Tissue-specific Multi-ISM attribution profiles across the 500 kb *ENPP2* locus on GTEx liver, brain, muscle and skin tracks. Colored shading corresponds to the four regulatory elements highlighted in (B) and (C). In gene tracks, orange and blue denote plus- and minus-strand genes. broadly shared promoter activity is tuned by tissue-specific enhancers driven by distinct regulatory programs.

We next asked what Multi-ISM attribution maps could reveal about the regulatory architecture of individual genes. For each gene, the full track-by-position Multi-ISM matrix provides a cross-tissue view of sequence attribution across the model input window. For example, the ribosomal protein gene *RPL5* exhibits a compact and largely uniform regulatory pattern, with influential nucleotides concentrated near the promoter and gene body, shared across tissues (**Supp. Fig. S6**). In contrast, *ENPP2*, which encodes the secreted lysophospholipase D autotaxin, displays a highly structured and tissue-specific regulatory landscape spanning nearly 200 kb (**Fig. 4B**). While its promoter was active in nearly all tracks, many distal regions were active in only a subset of tissues. We examined the *ENPP2* regulatory landscape more closely in four GTEx tissues in which the gene is expressed—liver, brain, muscle, and skin—and highlighted four candidate enhancers with tissue-specific Multi-ISM profiles (**Fig. 4B–D**). The blue-shaded enhancer, located 48.6 kb upstream of the *ENPP2* transcription start site (TSS), showed liver-specific attribution centered on a FOXA-family motif identified using tangermeme [18], consistent with the established role of FOXA factors in hepatic regulatory programs [19, 20]. The red-shaded enhancer, 14.0 kb upstream of the TSS, showed its strongest attribution in muscle and contained a MEF2D motif, consistent with the role of MEF2-family factors in muscle regulation [21, 22]. The green-shaded enhancer was most active in brain and contained a SOX-family motif, consistent with the central role of SOX factors in neural cell states [23, 24]. The purple-shaded enhancer, located 94.4 kb from the TSS, showed preferential activity in skin and contained an ETS-family motif matching ELF3, consistent with the function of ELF3 and related ETS factors in epithelial transcriptional regulation [25, 26]. An additional example of tissue-specific regulatory architecture at the *BMP7* locus is shown in **Supp. Fig. S6**. By mapping each gene’s regulatory landscape across long genomic windows and hundreds of tracks, Multi-ISM captures architectures ranging from compact, promoter-dominated regulation like *RPL5* to more complex patterns, such as those of *ENPP2* and *BMP7*, in which

### 3.5 Constrained genes show smaller predicted mutational effects and more diverse regulatory patterns

The track-by-position Multi-ISM matrix provides a gene-centered representation of the predicted regulatory landscape across both genomic position and cellular context. We therefore asked whether different summaries of this matrix could capture broader properties of gene regulation.

We first quantified a gene’s regulatory complexity using the effective rank of its track-by-position Multi-ISM matrix (**Methods**). Effective rank was derived from the entropy of the matrix’s singular-value spectrum and provides a relative, gene-level measure of regulatory complexity across both the track and genomic axes. Low effective rank indicates that the attribution matrix is dominated by a small number of regulatory patterns shared across tracks, whereas high effective rank indicates that distinct regulatory patterns contribute across tissue contexts.

Effective rank was positively associated with the GTEx tissue-specificity index *τ* (Spearman’s *R* = 0.63; **Fig. 5A**) [28], consistent with tissue-restricted genes drawing on more context-specific regulatory patterns than broadly expressed genes. Effective rank also correlated with the number of rE2G enhancer links per gene (Spearman’s *R* = 0.51; **Fig. 5A**; **Methods**) [29]. We next compared effective rank to two promoter properties measured by Tan et al. in an MPRA of enhancer– promoter combinations in K562 cells [27]. Basal activity, the expression a promoter drives when paired with inactive control enhancers, was negatively associated with effective rank (Spearman’s *R* = *−*0.51; **Fig. 5A**). Responsiveness, the slope relating a promoter’s expression gain to the activity of the enhancer paired with it, was positively associated with effective rank (Spearman’s *R* = 0.38; **Fig. 5A**). Genes with more complex regulatory architecture therefore tend to have weakly active promoters that gain more when paired with an active enhancer. Promoter responsiveness correlated more strongly with effective rank calculated within promoter (Spearman’s *R* = 0.44) and distal non-coding (Spearman’s *R* = 0.43) regions than with the full 500-kb window measure (**Supp. Fig. S7**).

**Figure 5:**
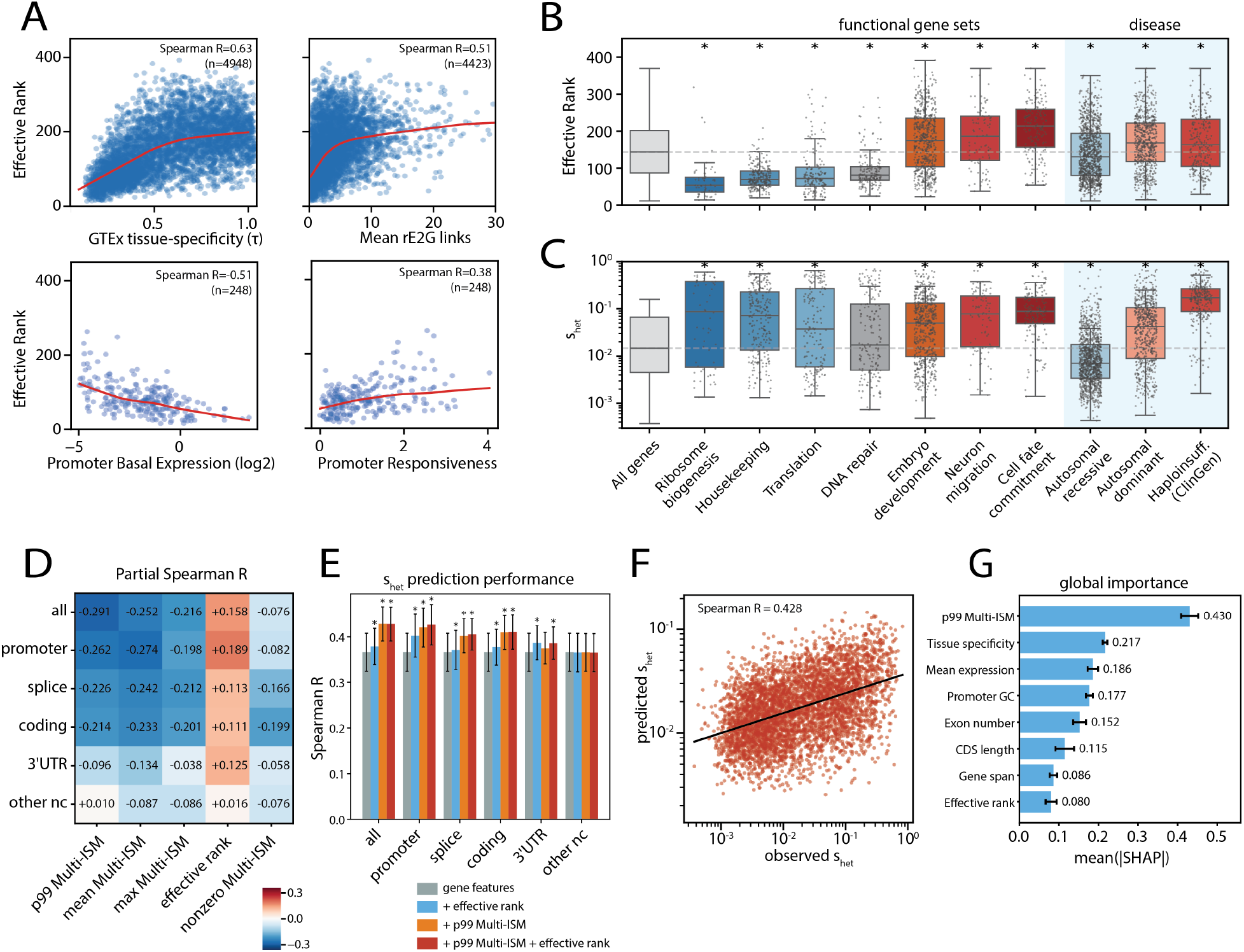
Multi-ISM captures gene regulatory complexity and mutational sensitivity. **A,** Effective rank versus GTEx tissue specificity (*τ*), mean ENCODE rE2G enhancer links across five cell lines, basal promoter activity, and promoter responsiveness measured by K562 MPRA [27]. Red lines, LOWESS fits. **B-C,** Effective rank (B) and gene constraint (*s*_het_; C) across functional and disease-associated gene sets. Boxes show median and interquartile range; whiskers extend to 1.5*×* the interquartile range. Gray points denote individual genes and are omitted for All genes. Dashed lines indicate the global median. Asterisks denote differences from All genes by two-sided Mann–Whitney U test with FDR correction (*q <* 0.001). **D,** Partial Spearman correlations between gene-level Multi-ISM summaries and *s*_het_, controlling for mean expression, tissue specificity, exon number, CDS length, gene span, and promoter GC content. Statistics were computed across the full 500-kb window (all) or within individual annotation categories. p99 Multi-ISM, 99th percentile of absolute non-zero effects; mean Multi-ISM, mean absolute non-zero Multi-ISM effect; max Multi-ISM, maximum absolute effect; effective rank, effective rank of the position-by-track matrix; nonzero Multi-ISM, fraction of non-zero matrix entries. **E,** Cross-validated prediction of *s*_het_ using gene features alone (gray), plus effective rank (blue), plus p99 Multi-ISM (orange), or plus both (red), evaluated separately by annotation category. Bars show Spearman correlation between pooled held-out predictions and observed *s*_het_; error bars show standard deviation across folds. Asterisks indicate improvement over gene features alone by one-sided Wilcoxon signed-rank test across folds (*P <* 0.05). **F,** Observed versus held-out predicted *s*_het_ for the full model using gene features, effective rank, and p99 Multi-ISM computed across the full 500-kb window. Predictions are pooled across held-out genes from all folds. **G,** Global feature importance in the full *s*_het_ prediction model, quantified by mean absolute SHAP value. Bars show mean importance across folds; error bars indicate the standard deviation.

By contrast, the total number of base pairs overlapping ENCODE candidate cis-regulatory elements within a 500-kb window was essentially uncorrelated with effective rank (Spearman’s *R* = *−*0.08; **Supp. Fig. S7**). Thus, effective rank does not simply reflect the abundance of nearby regulatory elements: a gene can reside within a broadly accessible regulatory neighborhood while still exhibiting relatively simple regulatory architecture.

We next compared effective rank across functional gene sets (**Fig. 5B**). Housekeeping and core cellular categories, including ribosome biogenesis, translation, and DNA repair, generally showed low rank. In contrast, developmental and cell-identity programs, including embryonic development, neuronal migration, and cell-fate commitment, showed higher effective rank. Gene sets at both ends of the effective-rank spectrum — ribosome biogenesis and translation at the low end, cell-fate commitment at the high end — had elevated constraint coefficients (*s*_het_) relative to all genes (**Fig. 5C**). Consistent with this observation, effective rank showed little marginal association with *s*_het_ across genes (Spearman’s *R* = 0.036).

We next asked whether effective rank or other Multi-ISM summaries were associated with constraint after controlling for mean expression, tissue specificity, exon number, CDS length, gene span, and promoter GC content (**Fig. 5D**). Across the full 500-kb window, the 99th-percentile absolute non-zero Multi-ISM effect (*p*99), which captures the upper tail of mutational effects while being robust to outliers, had the strongest partial correlation with *s*_het_ (partial Spearman’s *R* = *−*0.291). Thus, more strongly constrained genes have smaller predicted upper-tail mutational effects. The relationship held within promoter, splice-site, coding, and (to a lesser extent) 3’ UTR regions. After accounting for covariates, effective rank also showed a weak, positive partial correlation with *s*_het_ (partial Spearman’s *R* = 0.158); among genes matched for expression and gene structure, those with more complex regulatory architecture tended to be more constrained. Consistently, adding *p*99 to a linear model of the same gene features improved 10-fold cross-validated prediction of *s*_het_ in every annotation category except distal noncoding (**Fig. 5E**); for the full 500-kb window, held-out Spearman’s R increased from 0.366 to 0.428 (*R*^2^ from 0.127 to 0.188; **Fig. 5F, Supp. Fig. S7**). In the full predictive model, *p*99 was the most important feature by mean absolute SHAP value (**Fig. 5G**; **Supp. Fig. S7**).

Together, these analyses identify distinct gene-level properties captured by Multi-ISM maps. Effective rank reflects the complexity and context dependence of regulatory architecture, whereas upper-tail mutational effect magnitude reflects sensitivity to strong sequence perturbations. After controlling for expression and gene structure, more strongly constrained genes show attenuated predicted mutational effects and higher effective rank. One possible explanation is that strong selection can favor regulatory redundancy and other buffering mechanisms that reduce sensitivity to individual genetic perturbations.

### 3.6 Multi-ISM rare-variant burdens improve personalized expression prediction

Previous studies have shown that sequence-to-function models such as Enformer, Borzoi, and AlphaGenome poorly predict personal gene expression from genotypes [30–33]. Fine-tuning these models on cohorts with matched personal genomes and transcriptomes has not consistently outperformed linear models such as PrediXcan elastic net evaluated on held-out individuals at the same genes [34, 35].

Most personal-expression studies have focused on common variants because cohorts with paired genotype and expression data remain relatively small, limiting the power to learn rare-variant effects. Nevertheless, sequence-based models may be most useful for rare variants with large regulatory effects. Borzoi distinguishes under- from over-expression outliers in GTEx zero-shot (AUROC 0.85) [36]. Consistently, a recent study found that fine-tuned Borzoi modestly outperformed elastic net for predicting protein abundance, with its advantage increasing at larger sample sizes and depending partly on rare variants [37].

Motivated by these observations, we hypothesized that sequence-to-function predictions could improve personalized gene-expression prediction by capturing the effects of variants too infrequent for an elastic net to learn reliably, without fine-tuning on personal genomes. Instead of modeling these variants individually in a PrediXcan-style framework, we used sequence-to-function predictions to aggregate their effects into a single burden score, which was then included as an additional feature in the elastic-net model (**Fig. 6A**). Multi-ISM efficiently enables this approach by pre-computing the effects of all single-nucleotide substitutions within a 500-kb window centered on each gene. For each individual, we calculated a Multi-ISM burden by summing the predicted effects of variants with MAF *<* 5% separately on each haplotype and averaging the two sums (**Methods**). We refer to the elastic-net model trained directly on common-variant genotype features (MAF *≥* 5%) as the *common-variant elastic net* (labeled in figure as EN-common), and to the model additionally including the Multi-ISM burden from variants with MAF *<* 5% as the *Multi-ISM–augmented elastic net* (labeled in figure as EN-common+MISM).

**Figure 6:**
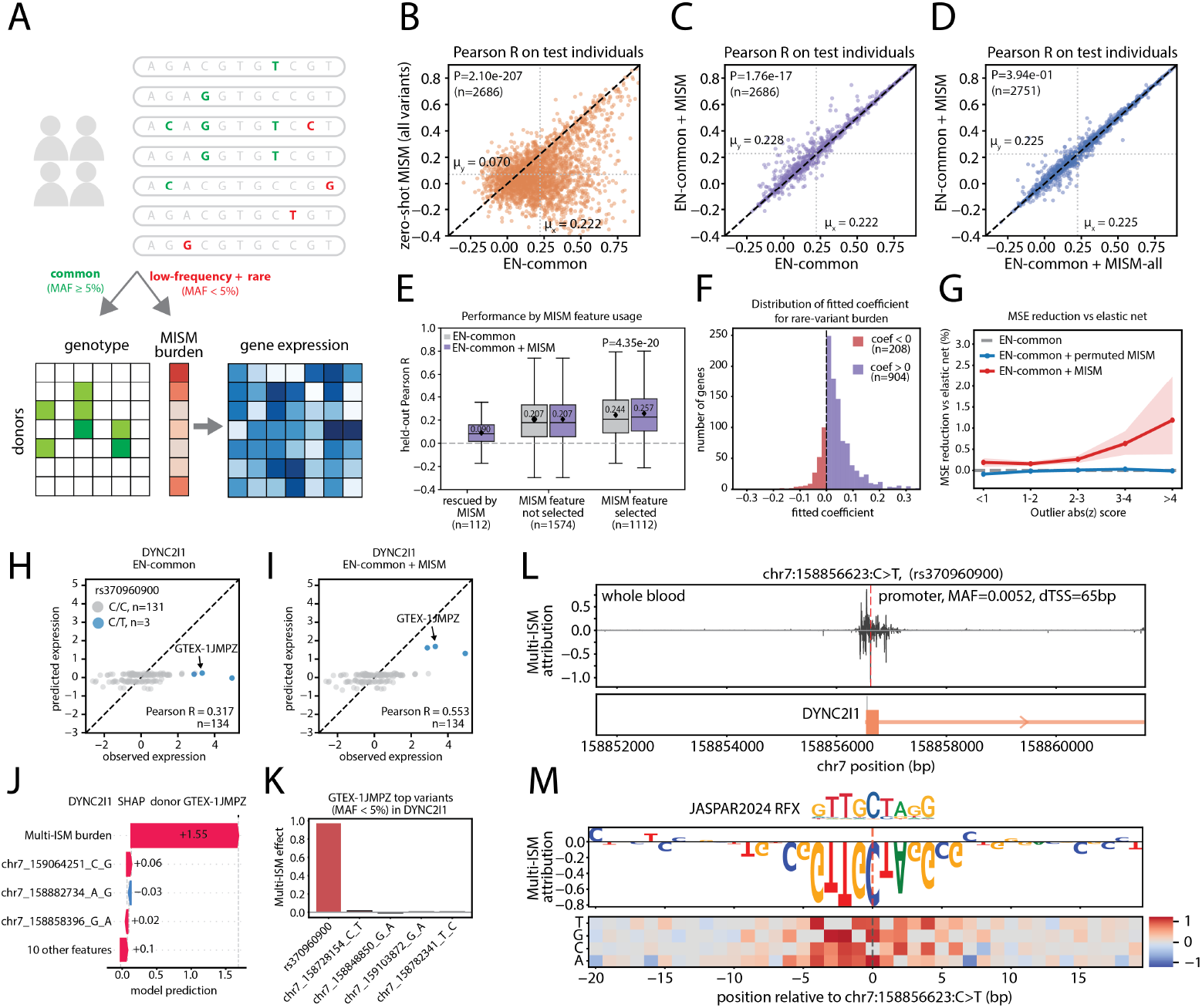
Multi-ISM–augmented elastic net improves personalized expression prediction. **A,** Schematic of the prediction framework. The *Multi-ISM–augmented elastic-net* model is trained using common-variant genotype features (MAF *≥* 5%) together with a burden feature obtained by aggregating Multi-ISM predictions for variants with MAF *<* 5%. Multi-ISM is abbreviated as MISM in this figure. **B,** Comparison of personalized gene-expression prediction by the *common-variant elastic net* (EN-common) and zero-shot Multi-ISM. EN-common uses common-variant genotypes, whereas zero-shot Multi-ISM predicts expression by summing the Multi-ISM effects of all variants carried by each individual within 500-kb window. Genes with defined correlations in both models (2686) are shown. **C,** Comparison of the EN-common with the *Multi-ISM–augmented elastic net* (EN-common + MISM), which adds the Multi-ISM burden aggregated from variants with MAF *<* 5% as an additional feature. *P* value is from a one-sided Wilcoxon signed-rank test. Genes with defined correlations in both models (2686) are shown. **D,** Comparison of EN-common augmented with either the Multi-ISM burden from variants with MAF *<* 5% (EN-common + MISM) or across all variants (EN-common + MISM-all). Genes with defined correlations in both models (2751) are shown. **E,** Held-out Pearson R stratified by Multi-ISM feature usage: genes rescued from a constant EN-common prediction (n=112), genes for which the Multi-ISM feature is not selected (n=1,574), and genes for which it is selected (n=1,112). One-sided Wilcoxon signed-rank test. **F,** Distribution of fitted Multi-ISM burden coefficients among genes for which the feature was selected; 904 were positive and 208 negative. **G,** Percentage reduction in mean squared error (MSE) relative to EN-common (dashed line), stratified by expression outlier score. EN-common + MISM is shown in red and a permuted-MISM control in blue. Shaded bands are 95% confidence intervals from 1000 bootstrap resamples. Outlier score is the absolute deviation from mean after regressing out covariates and the lead eQTL (Methods). **H-I,** Observed versus predicted *DYNC2I1* whole blood expression in held-out individuals using H, EN-common or I, EN-common + MISM. Blue points are heterozygous carriers of rs370960900 C*>*T (MAF=0.0052); gray points are C/C homozygotes. Carrier GTEX-1JMPZ is highlighted. **J,** SHAP waterfall plot of the EN-common + MISM model prediction for donor GTEX-1JMPZ. The Multi-ISM burden is the dominant feature (SHAP value = 1.55). **K,** Multi-ISM-predicted effects of top scoring variants carried by GTEX-1JMPZ within the 500-kb *DYNC2I1* regulatory window. **L,** Multi-ISM attribution around rs370960900 (chr7:158,856,623 C*>*T) for GTEx whole-blood track. The variant lies in the *DYNC2I1* promoter. **M,** Multi-ISM substitution map around rs370960900, shown as a reference sequence logo (top) and predicted effect of all three substitutions at each position (bottom). Downward letters and pos^1^it^4^ive substitution effects across the RFX motif indicate that mutating the motif increases predicted expression, consistent with a repressive site. the remaining 1,251 genes, all coefficients were shrunk to zero, leaving constant predictions. Among the 2,686 genes with defined elastic-net correlations, directly aggregating Multi-ISM scores across variants of all allele frequencies achieved a mean Pearson’s R of 0.070 (Fig. 6B), consistent with previous studies showing that zero-shot sequence-to-function models underperform elastic net for this task [30–33].

We evaluated personalized gene-expression prediction using GTEx v8 genotypes and covariate-corrected whole-blood expression from 670 donors (**Methods**). Among the 3,937 genes with both Multi-ISM scores and whole-blood expression data, the *common-variant elastic net* produced non-constant predictions for 2,686 genes, with a mean Pearson’s R of 0.222 in held-out individuals; for

We next tested whether incorporating the Multi-ISM burden could improve prediction beyond the *common-variant elastic net*. For each gene and individual, we aggregated the predicted effects of variants with MAF *<* 5% into a single burden and included it as an additional feature during elastic-net training (**Fig. 6A**). The resulting *Multi-ISM–augmented elastic net* improved the mean Pearson’s R in held-out individuals from 0.222 to 0.228 (*P* = 1.76 *×* 10*^−^*^17^, one-sided Wilcoxon signed-rank test; **Fig. 6C**). It also outperformed a baseline model trained with permuted Multi-ISM burden (**Supp. Fig. S8**).

Replacing the Multi-ISM burden from variants with MAF *<* 5% with a burden aggregated across variants of all allele frequencies did not further improve performance, indicating that the predictive gain was driven primarily by low-frequency and rare variants (MAF *<* 5%; **Fig. 6D**; **Supp. Fig. S8**). To test whether this benefit simply reflected the absence of variants with MAF *<* 5% from the baseline elastic net, we progressively expanded the genotype feature set of the baseline model to variants with MAF *≥* 1%, and then to all variants without allele-frequency filtering. Adding the same Multi-ISM burden continued to improve prediction in each setting, indicating that its benefit was not simply due to the exclusion of lower-frequency and rare variants from the baseline elastic-net model (**Supp. Fig. S8**).

Although the overall improvement was modest, stratifying genes by how the Multi-ISM burden feature was used revealed that the gains were concentrated in a subset of genes. Of the 1,251 genes for which the *common-variant elastic net* shrank every coefficient to zero, adding the Multi-ISM burden produced a non-constant prediction for 112. For 1,574 genes, the Multi-ISM burden was not selected, and the two models were effectively identical (mean Pearson’s *R* = 0.207 for both). In contrast, on the 1,112 genes in which the Multi-ISM burden was selected as a feature, the *Multi-ISM–augmented elastic net* outperformed the common-variant model (mean Pearson’s *R* = 0.257 versus 0.244; *P* = 4.35 *×* 10*^−^*^20^, one-sided Wilcoxon signed-rank test; **Fig. 6E**). Notably, the learned coefficient for the Multi-ISM burden was positive for 904 of these genes (81%; **Fig. 6F**), indicating that the direction of the burden as predicted by Multi-ISM was generally concordant with personalized gene expression.

We next compared the two models across individual gene-by-donor predictions. The relative reduction in prediction error was greatest at the gene–donor pairs with the largest expression-outlier scores (**Fig. 6G**; **Methods**). Together, these results are consistent with the *Multi-ISM–augmented elastic net* improving personalized expression prediction by capturing high-effect rare variants that contribute to outlier expression.

To illustrate how Multi-ISM burden can improve personalized expression prediction, we examined *DYNC2I1*, an OMIM gene for which held-out performance increased from Pearson’s *R* = 0.317 with the *common-variant elastic net* to *R* = 0.553 with the *Multi-ISM–augmented elastic net*. The gain was driven largely by three individuals with markedly elevated *DYNC2I1* expression, all heterozygous for the rare rs370960900 C*>*T allele (MAF = 0.0052), whereas the remaining 131 test individuals were C/C homozygotes (**Fig. 6H–I**). The *common-variant elastic net* systematically under-predicted expression in these carriers, while adding the Multi-ISM burden substantially improved agreement with the observed values.

For a representative carrier GTEX-1JMPZ, SHAP analysis identified the Multi-ISM feature as the dominant contributor to the prediction (SHAP value =1.55), with rs370960900 accounting for most of the burden (**Fig. 6J–K**). Multi-ISM predicted that this C*>*T promoter variant would increase whole-blood *DYNC2I1* expression by log2FC = 0.973 on the carrying haplotype, approximately doubling its predicted transcript abundance (**Fig. 6K**). Consistent with this prediction, PromoterAI also assigned the variant a strong expression-increasing effect (score = 0.82). The variant falls within an RFX motif, and mutations across that motif are uniformly predicted to be expression-increasing, suggesting that disruption of a repressive RFX site could account for the elevated expression (**Fig. 6L–M**).

Multi-ISM also leverages low-frequency and rare variants outside promoters to improve personalized expression prediction (**Supp. Fig. S9**). For *MAN1B1*, Multi-ISM prioritized a rare variant (rs115944822, chr9:137089241 A*>*G; MAF = 0.0030) annotated in ClinVar as creating a cryptic splice acceptor (accession: VCV004354633.1), and predicted the increased expression observed in two carriers. For *PRPH2*, Multi-ISM prioritized a rare 3*^′^*-UTR variant (rs73426405, chr6:42697456 C*>*T; MAF = 0.0090) and correctly predicted increased *PRPH2* expression in carriers. Together, these examples illustrate how incorporating Multi-ISM rare-variant burdens can improve PrediXcan-style personalized gene-expression prediction by capturing regulatory effects that are difficult to learn directly from cohort data.

## 4 Discussion

Here, we introduce Multi-ISM, an architecture-agnostic framework that combines sparse recovery with adaptive pruning to perform efficient saturation mutagenesis across the large input windows of modern sequence-to-function models. By measuring multiple spatially separated mutations per sequence and adaptively refining their estimated effects, Multi-ISM generates 500-kb mutational maps using 45-fold fewer model evaluations than exhaustive single-variant ISM, while matching, and in several benchmarks exceeding, its variant-effect prediction accuracy.

Comprehensive mutational atlases catalog the regulatory effects learned by sequence-to-function models [3]. However, generating such maps by brute-force single-variant scoring requires millions of model evaluations per gene, and a single pre-computed map reflects the particular capabilities and biological scope of its underlying model, including its training data, predicted molecular readouts, and cellular contexts. New mutational maps are needed as these capabilities expand. Comprehensive mutational mapping is therefore a recurring computational challenge in interpreting evolving sequence-to-function models.

Multi-ISM addresses this challenge because it operates only through model inputs and outputs and does not depend on internal model architecture. We demonstrate portability across transformer- and state-space-based models, including Flashzoi, Borzoi-Prime, and Cerberus, spanning different mutational windows, molecular assays, and cellular contexts. Applied to Cerberus, Multi-ISM extended mapping to cell types that other models do not predict; for a fine-mapped *RASGEF1C* microglia eQTL, it predicted a strong microglial effect largely absent from other cell types and bulk-tissue tracks. Multi-ISM also generalizes across readouts; from the same model evaluations, we derive mutational effects on splice-junction usage and polyadenylation-site usage in addition to total transcript abundance. Together, these properties make Multi-ISM a scalable procedure for generating comprehensive mutational maps as sequence-to-function models expand to new architectures, functional measurements, and cellular contexts.

We applied Multi-ISM to 5,000 genes, including 3,317 OMIM genes, generating gene-centered attribution maps of predicted regulatory effects across genomic positions and cellular contexts. The maps are available at https://multi.ism.research.calicolabs.com/. Beyond individual variants and regulatory elements, these maps yield complementary gene-level summaries. Effective rank captures the diversity of predicted regulatory patterns across positions and tracks and correlates with tissue specificity, enhancer connectivity, and promoter responsiveness. The upper tail of absolute variant effects (*p*99) summarizes predicted sensitivity to individual mutations. After accounting for expression, tissue specificity, and gene structure, more strongly constrained genes had smaller predicted upper-tail effects and modestly higher effective rank, a pattern consistent with regulatory buffering [38, 39]. Large-scale, gene-centered saturation mutagenesis maps enable systematic comparisons of predicted regulatory properties across genes.

Personalized gene-expression prediction remains a major challenge for sequence-to-function models. Zero-shot predictions of these models explain substantially less cross-individual expression variation than cohort-trained linear models, and fine-tuning with matched personal genomes and transcriptomes yields only partial improvements [30–32, 34, 35, 40]. For fine-mapped eQTLs, sign prediction by Borzoi and AlphaGenome approaches chance for small predicted effects but is most accurate for large ones (**Supp. Fig. S1**), consistent with recent benchmarking [41]. We therefore asked whether sequence-model predictions could complement cohort-trained predictors by capturing rare variant effects that are difficult to learn directly from cohort data. Aggregating pre-computed Multi-ISM effects into a gene-level burden modestly but significantly improved held-out common-variant elastic-net predictions without fine-tuning the underlying sequence model. The gain was concentrated at gene–individual pairs with outlier expression, consistent with rare variants having large effects in only a small subset of gene–individual pairs [36, 42, 43]. In the examples we examined, the prioritized variants lay in promoters, introns, cryptic splice sites, and 3*^′^*-UTRs, spanning transcriptional and post-transcriptional regulation.

Multi-ISM relies on an assumption of regulatory sparsity, whereby only a small fraction of the possible substitutions within a 500 kb window are expected to have predictable effects on the target output. Under this assumption, sparse recovery reduces the number of measurements required and produces compact attribution maps that can be stored and queried efficiently. Because regulatory effects become increasingly sparse with distance from the target gene, expanding the mutagenized window adds relatively few additional nonzero effects. Consistent with this, the performance gap between 100-kb and 500-kb windows for Flashzoi nearly closed as measurements increased, while Cerberus achieved comparable performance over a 700-kb window at the same budget. Yuzu [10] also uses sparse recovery, but exploits sparse activation differences in convolutional layers, making its acceleration architecture dependent; Multi-ISM assumes sparsity in the variant-effect landscape itself, trading exact recovery for architecture independence. This approach can shrink or omit genuine weak effects through *L*_1_ regularization.. Multi-ISM should therefore be interpreted as recovering the most stable effects supported by a given measurement budget, a reasonable trade-off given that the eQTL benchmark revealed the zeroed effects are less reliable.

A second key assumption of Multi-ISM is that the effects of spatially separated mutations are approximately additive. This assumption suits our goal, which is to estimate marginal single-variant effects rather than higher-order interactions. We reduce exposure to local epistasis by enforcing a minimum distance between mutations introduced into the same sequence. Under this distance-constrained design, adding a global nonlinear mapping to an additive genotype–phenotype model with MAVE-NN did not improve held-out prediction (**Methods, Supp. Fig. S2**) [11, 44]. Interactions between distant mutations, such as in an enhancer and its target promoter, escape the spacing constraint and could reduce recovery accuracy. When epistasis itself is the quantity of interest, however, frameworks designed specifically to model sequence interactions would be more appropriate.

Finally, extending Multi-ISM to indels and structural variants could be particularly valuable given their substantial contribution to disease [43], but remains challenging. Sequence-to-function predictions for these variant classes are less robust [45], and the mutation space grows rapidly with variant length. Future work could therefore explore alternative mutation encodings and more efficient search strategies to extend Multi-ISM beyond single-nucleotide variants.

## 5 Methods

### 5.1 Multi-ISM

Multi-ISM is a multi-stage framework for efficient in silico saturation mutagenesis that combines (i) a sparse linear formulation of variant effect estimates, (ii) a structured mutation design that enforces minimum spacing between variants, and (iii) an iterative adaptive pruning strategy to improve efficiency and accuracy of estimation. Together, these components enable scalable estimation of variant effects with fewer model evaluations, while reducing dependence on a single reference background.

#### 5.1.1 Sparse recovery formulation

To improve the efficiency and robustness of in silico saturation mutagenesis, Multi-ISM introduces multiple variants per forward pass while enforcing a minimum spacing constraint between mutated positions. Each forward pass measures the aggregate output of multiple variants, substantially reducing the number of forward passes compared to single-variant ISM. Because each variant appears across many sequence backgrounds, its estimated effect averages over those contexts and can differ from its effect in the reference sequence.

We approximate each forward pass output as the sum of the individual variant effects at the mutated positions included in that pass. Formally, let:

- *y ∈* ℝ*^n^*: vector of measurements (model outputs from *n* forward passes),
- *x ∈* ℝ*^p^*: vector of single-variant effects, where *p* = 3*L* for a sequence of length *L*,
- *A* ∈ {0, 1}*^n×p^*: design matrix, where *A_ij_* = 1 if variant *j* is included in forward pass *i*, and 0 otherwise.

Under an additive approximation, the full set of observations can be written compactly as

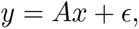

where *ɛ* captures model stochasticity, numerical error, and residual non-additive effects.

This formulation relies on two assumptions: (1) variant effects are approximately additive when variants are separated by at least a specified distance, such that higher-order interactions are rare; and (2) within the long context mutation window around a gene, only a small subset of variants have non-negligible regulatory effects, implying that *x* is sparse with *s ≪ p* non-zero entries.

Compressed sensing theory provides guidance on the number of measurements required to recover such a signal. For random measurement designs meeting sparse-recovery conditions, an *s*-sparse signal can be recovered with a measurement count of order

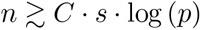

measurements, where *C* depends on the design and noise level. Under these conditions, measurement requirements grow primarily with the number of non-zero effects and only logarithmically with the number of candidate variants. This scaling motivates the sparse-recovery approach used by Multi-ISM. Randomized mutation assignment varies the combinations in which variants are evaluated, while the minimum spacing constraint is intended to preserve approximate additivity.

We estimate *x* by solving the *L*_1_-regularized least-squares problem

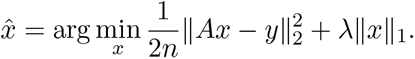

which promotes sparse solutions and provides stable recovery in the presence of noise and approximation error. We used the `celer` package to solve the LASSO problem and selected *λ* = 10*^−^*^4^ by 5-fold cross-validation over *λ ∈* [10*^−^*^7^, 10*^−^*^6^*,…,* 10^2^] [46].

#### 5.1.2 Mutation matrix design

Let **s** = (*s*_1_*, s*_2_*,…, s_L_*) denote a sequence of length *L*, where *s_i_ ∈* {A, C, G, T}. Given a minimum distance constraint *d*, we partition the sequence into non-overlapping bins

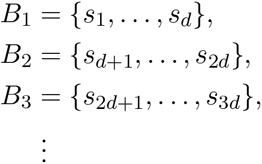

with the final bin truncated if necessary. Each bin corresponds to a contiguous subsequence of length *d*.

We consider all single-nucleotide substitutions within each bin, yielding 3*d* possible mutations per bin. Mutation sequences are generated by alternating between odd and even bins. In one forward pass, we include one mutation from each odd bin (*B*_1_*, B*_3_*, B*_5_*,…*); in the next, we include one mutation from each even bin (*B*_2_*, B*_4_*, B*_6_*,…*). This alternating procedure continues until each mutation has been included exactly *k* times. The total number of forward passes constructed is *n* = *d* (positions per bin) *×* 3 (mutations per position) *× k* (repetition) *×* 2 (odd/even bins) = 6*dk,* which depends only on the distance constraint *d* and the mutation repetition count *k*, and is independent of the total sequence length *L*. In our study, we fix *L* = 500 kb, *d* = 50 bp.

We call this the **base Multi-ISM** design. This design ensures that (i) all mutations within a forward pass are separated by at least *d*, enforcing the distance constraint required for approximate additivity; (ii) mutations are randomly assigned across forward passes, reducing correlations between columns of the design matrix *A*; and (iii) each mutation is evaluated across multiple sequence backgrounds.

#### 5.1.3 Response vector construction

For a given mutation design vector, row of *A*, we construct the corresponding mutated sequence and evaluate the sequence model in a single forward pass, using the forward strand or its reverse complement with equal probability. For computational efficiency, Multi-ISM model evaluations were performed using Flashzoi [14], an optimized Borzoi-derived implementation that replaces the original transformer attention with a FlashAttention-compatible architecture while retaining the Borzoi prediction framework.

For each target gene, we aggregate predicted coverage over the set of genomic bins *G* = {*g*_1_*,…, g_M_*} overlapping the gene’s exons. The response *y_i_* for the *i*-th forward pass is defined as the log fold-change between the alternate and reference alleles:

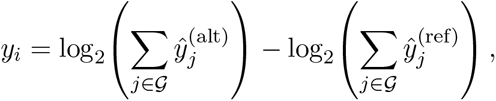

where 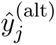 and 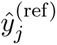 denote the model-predicted coverage for bin *j* under the alternate and reference sequences, respectively. Alternate sequences are evaluated in randomly selected strand orientations, and each response is computed against the cached reference prediction in the same orientation.

#### 5.1.4 Multi-ISM with adaptive pruning

In practice, regulatory variant effects exhibit a highly skewed distribution, with a small number of variants dominating the signal. Although sparse regression methods such as LASSO readily identify these dominant variants, even small estimation errors in their effects can induce large residuals in the response vector that obscure moderate-effect variants. As a result, substantially more measurements are required to resolve moderate-effect variants accurately.

To address this issue, we introduce **adaptive Multi-ISM**, an iterative pruning strategy built on top of the base Multi-ISM framework. The key idea is to progressively identify high-effect regulatory variants, estimate their effects accurately, and remove their contribution from the response vector, to allow moderate-effect variants to be recovered with improved stability and sensitivity.

This coarse-to-fine recovery strategy substantially improves overall recovery efficiency and forms the final variant effect estimation pipeline used in our analyses.

##### Initial iteration

In the initial iteration, we perform base Multi-ISM across the full 500 kb target region as described above. Each mutation is repeated 20 times (*k* = 20), resulting in a total of 6,000 forward passes, which provides a coarse but comprehensive estimate of the regulatory landscape.

##### Subsequent iterations

In each subsequent iteration, we apply the following three steps:

- **High-effect variant pruning.** Variants are ranked by the absolute magnitude of their estimated effects from the previous iteration, and the top-ranking positions (default: 200) are selected as high-effect sites. For each selected position, all three possible nucleotide substitutions are re-measured using background-augmented single-ISM. Specifically, each substitution is evaluated in randomly mutated sequence backgrounds (default: 4), using both strand orientations. Its effect is calculated relative to the matched background without that substitution and averaged across backgrounds and orientations to improve robustness [11]. The selected positions are then added to a cumulative set of **resolved variants** and excluded from subsequent regression designs. This step requires 200 *×* 3 *×* 4 *×* 2 = 4,800 forward passes, corresponding to 200 positions, three substitutions per position, four sequence backgrounds, and two strand orientations.
- **Mutation design.** A new base Multi-ISM design is generated while excluding all previously resolved variants. This ensures that new forward passes focus exclusively on unresolved genomic positions. In this stage, we set *k* = 34, corresponding to 10,200 forward passes.
- **Estimate variant effects.** Mutation design matrices and response vectors from all iterations are concatenated. Effects of resolved variants are removed from the response, and the corresponding columns are excluded from the design matrix. LASSO regression is then applied to estimate the effects of the remaining variants.

Overall, this adaptive pruning strategy improves the conditioning of the sparse regression problem by isolating and removing dominant variant effects early. This enables subsequent regression steps to more effectively resolve moderate-effect regulatory variants, resulting in more stable and sensitive recovery of the regulatory landscape. In practice, five iterations, 66,000 forward passes (4800 + 10200) *×* 4 + 6000, are typically sufficient to produce high-quality variant effect estimates for a 500-kb window of a gene across all tasks.

### 5.2 Genome-scale Multi-ISM resource

We applied adaptive Multi-ISM to generate nucleotide-resolution variant-effect maps for 5,000 protein-coding genes across 500 kb gene-centered windows. The gene set combined loci used in benchmarking and downstream analyses with a broader collection of disease-relevant and regulatory genes, including 3,317 Mendelian disease genes curated from OMIM by Minikel et al. [47]. The OMIM set was restricted to established single-gene disease associations, mapped to current gene symbols, and limited to protein-coding genes in GENCODE v38. Each gene was processed independently, and the resulting Multi-ISM maps were used as the shared resource for the analyses in this study.

### 5.3 Sensitivity analysis

We characterized the sensitivity of base Multi-ISM to key design parameters at the GATA1 locus. Reconstruction accuracy was evaluated by Pearson correlation with exhaustive single-mutation ISM in the central 10-kb window.

#### Minimum distance between mutations

To evaluate the minimum spacing constraint, we fixed the context window at 100 kb and the total number of measurements at 15,000, and tested effect of *d ∈ {*10, 20, 50, 100*}* bp. Smaller values of *d* include more mutations per sequence and therefore provide more repeated observations per position, but at the cost of reduced independence between nearby mutations. We found that *d* = 10 and *d* = 20 showed worse performance despite the higher per-position sampling rate, consistent with greater departures from additivity when mutations are closely spaced (**Supp. Fig. S2**). In contrast, *d* = 50 and *d* = 100 performed comparably well. We selected *d* = 50 bp as the default spacing constraint.

#### Mutation window size

We fixed *d* = 50 bp and asked Multi-ISM to recover variants effect on 100-kb and 500-kb windows across measurement budgets ranging from 15,000 to 300,000. At 15,000 measurements, Multi-ISM with the 100-kb window substantially outperformed the 500-kb window, as the latter attempts to estimate five times as many variant effects from the same number of observations. But this gap closed with increasing data, and at 250,000 measurements, both configurations achieved nearly identical correlation (**Supp. Fig. S2**). This observation is consistent with compressed sensing theory, which predicts that, for suitable measurement designs, the number of measurements required for sparse recovery scales linearly with the number of truly non-zero signals and only logarithmically with the total dimensionality of the problem. Although expanding the window from 100 kb to 500 kb increases the number of variant positions to estimate by approximately five-fold, regulatory effects are very sparse further away from gene, so the number of non-zero coefficients grows by only a small fraction. Consequently, the larger window requires only modestly more measurements to achieve equivalent reconstruction accuracy.

### 5.4 Multi-ISM extension to additional architectures

#### Borzoi-Prime

Borzoi-Prime is a transformer-based sequence-to-function model that takes 524,288 bp of input sequence and predicts binned coverage tracks at 32 bp resolution for human and mouse functional tracks including ATAC, DNase, CAGE, RNA and scRNA [15]. We applied the same adaptive Multi-ISM pipeline to the Borzoi-Prime model (human head, fold 0, the fold with best GTEx eQTL performance), using a 500-kb mutational window centered on each gene and the same hyperparameters as the Flashzoi runs (random reverse-complement augmentation, bfloat16 precision, and five adaptive pruning iterations). The only difference lay in the model and its output space: Borzoi-Prime predicts 5,432 tracks, from which we collected variant effect scores on the 2,444 CAGE, RNA-seq, and scRNA-seq expression tracks. The Borzoi-Prime model is available at https://github.com/calico/baskerville.

#### Cerberus

Cerberus is a long-range sequence-to-function model that takes 786,432 bp of input sequence and predicts binned coverage tracks at 32 bp resolution for human and mouse functional tracks including ATAC, DNase, CAGE, RNA and scRNA [16]. The trunk comprises a DNA convolution stem, convolutional layers, and Hydra (bi-directional Mamba) blocks, followed by output heads. We applied the same adaptive Multi-ISM pipeline to the Cerberus model (human head, fold 6, the fold with best GTEx eQTL performance), keeping most hyperparameters shared with the Flashzoi and Borzoi-Prime runs (random reverse-complement augmentation, bfloat16 precision, and five adaptive pruning iterations). Two features distinguished the Cerberus run: we used a larger 700-kb mutational window centered on each gene, and the model predicts 8,361 tracks, from which we collected variant effect scores on the 3,171 CAGE, RNA-seq, and scRNA-seq expression tracks. The Cerberus model is available at https://github.com/calico/baskerville.

### 5.5 Multi-ISM extension to additional readouts

The standard Multi-ISM response is derived from a scalar expression readout of each model prediction: predicted coverage is summed across exonic bins and log-transformed [1]. Then, difference between mutant and reference sequences is used as the response for Multi-ISM regression to estimate effect of individual variant. The expression readout primarily captures changes in overall transcript abundance and may be insensitive to transcript-structure changes, such as splicing, or alternative polyadenylation.

Multi-ISM can similarly be applied to other scalar readouts derived from the same model output, provided that the effects of spatially separated variants on the readout are approximately additive. We therefore defined two additional readouts that quantify local transcript structure from the model outputs. All readouts are computed from the same model forward passes and therefore require no additional model evaluations.

#### Splice-junction usage (SJ usage)

To quantify splice-site usage, we calculate the log_2_ ratio of exon-side to intron-side coverage at each exon–intron junction. Coverage is aggregated over adjacent, length-matched windows of up to five output bins on each side of the boundary. This produces one SJ-usage readout per splice site and target. Splice sites are defined from GENCODE v38 exon–intron junctions.

#### Polyadenylation site usage (PAS usage)

To quantify alternative polyadenylation usage, we calculate the log_2_ ratio of distal to proximal PAS coverage, with coverage at each PAS summed over the four output bins immediately upstream of and including the PAS bin. For each gene, the 3*^′^*-most PAS is used as the fixed distal site and paired with each proximal PAS at least 128 bp upstream, yielding one PAS-usage readout per PAS pair and target. PASs are derived from PolyA-DB v3, restricted to sites with sufficient 3*^′^*-reads support, no antisense-gene overlap, and linkage to GENCODE v38 annotations.

#### Variant-effect estimation

Variant effects on each readout are estimated in the same manner as for the standard expression readout. For each Multi-ISM sequence, we calculate the change in the corresponding readout between the multi-mutant and reference sequences (ΔSJ-usage or ΔPAS-usage), and use these differences as the response vector for Multi-ISM regression. The resulting regression coefficients therefore estimate the effect of each individual nucleotide substitution on the corresponding readout. Regression is performed independently for each splice site or PAS pair. For SJ-usage, only variants within *±*5 kb of the splice site are included. For PAS-usage, variants are restricted to the genomic window [min(prox, dist) *−* 10 kb, max(prox, dist) + 10 kb].

### 5.6 Comparison with SQUID/MAVE-NN

SQUID is an interpretation framework that generates in silico mutagenesis libraries and fits surrogate models to DNN predictions [11, 44]. Although SQUID also evaluates multi-mutant sequences, it is designed for dense local surrogate modeling, typically using large in silico MAVE libraries over short sequence windows (100,000 measurements for hundreds of bp). Its surrogate model MAVE-NN can account for global nonlinearity and heteroskedastic noise. We therefore benchmarked MAVE-NN against the linear solver used by Multi-ISM to evaluate whether these additional modeling components improve variant-effect recovery under the Multi-ISM mutation design.

We compared MAVE-NN with celer elastic-net regression, which includes the LASSO used by Multi-ISM as the special case *ℓ*_1_ ratio = 1, on identical Multi-ISM-generated measurements at the GATA1 locus, with a minimum distance of *d* = 50 bp between mutations and focusing on a single K562 RNA-seq track ENCFF278EZT. MAVE-NN was fit with an additive genotype–phenotype map, global epistasis regression with 100 hidden nodes, and a heteroskedastic skewed-*t* noise model, as recommended by its authors. For each experiment, we generated 30,000 mutated sequences and split them into training, validation, and test sets at an 8:1:1 ratio. We tuned regularization strength *θ* of MAVE-NN, *ℓ*_1_ ratio and *α* of elastic net on the validation set.

For 1 kb sequences, simple elastic net (*ℓ*_1_ ratio = 0.5, and *α* = 10*^−^*^6^) and MAVE-NN (*θ* = 0.1) achieved the same correlation with measurement on hold-out test set (Pearson’s R=0.986), showing no improvement in held-out correlation from the global nonlinear mapping and noise model in this setting (**Supp. Fig. S2**). For 10 kb sequences, LASSO (*ℓ*_1_ ratio = 1 and *α* = 10*^−^*^5^) achieved Pearson R=0.963 on the test sequences, whereas MAVE-NN (*θ* = 10*^−^*^4^) achieved Pearson R=0.881, suggesting that as sequence size increases, LASSO better models the high-dimensional sparse signal, rather than the ridge-regularized additive map used by MAVE-NN (**Supp. Fig. S2**).

MAVE-NN also does not scale to the long-context setting targeted by Multi-ISM. In our experiments, fitting MAVE-NN required 7.52 GB of GPU memory for 10 kb inputs, which projects to roughly 400 GB for 500 kb inputs for a single output task. This memory burden is likely due to the dense gradient update implemented in MAVE-NN.

### 5.7 Agreement between Multi-ISM and single-ISM

Throughout the manuscript, single-ISM refers to measuring the effect of one variant at a time. Unless otherwise specified, we use Flashzoi fold 0; comparisons involving Borzoi-Prime or Cerberus use the corresponding model and fold described above. The input sequence is centered on the midpoint of the target gene, and both reference and alternate sequences are evaluated with reverse-complement augmentation. Predicted coverage is aggregated over output bins overlapping the gene’s exons, and the variant effect is defined as the log fold-change between the alternate and reference alleles.

Because the reference sequence needs to be evaluated only once for a given gene, we exclude this shared reference evaluation when estimating computational cost. Exhaustive single-ISM across a 500-kb mutagenesis window therefore requires approximately 500,000 *×* 3 *×* 2 = 3,000,000 forward passes, corresponding to three possible nucleotide substitutions per position and reverse-complement augmentation.

For comparisons between Multi-ISM and single-ISM on selected genes, Multi-ISM was performed across the full 500-kb mutagenesis window, whereas single-ISM was restricted to a 10-kb window centered on the midpoint of the gene body because of its substantially greater computational cost. Correlation between the two methods was therefore evaluated within this 10-kb window, restricted to variants with an absolute single-ISM effect greater than 0.01 to focus on effects more reliably estimated by single-ISM.

### 5.8 CAGI5 MPRA Saturation Mutagenesis Benchmark

To benchmark Multi-ISM performance, we used the CAGI5 MPRA datasets previously analyzed in the Borzoi study [1, 48]. Of the 14 CREs evaluated in Borzoi, we restricted our analysis to CREs satisfying the following criteria: (1) located within Multi-ISM context window of 250 kb from midpoint of gene body; (2) containing at least 300 variants with measured absolute logFC *>* 0.1; and (3) at least 20 variants with Flashzoi single-ISM predicted absolute logFC effect *>* 0.1. These filters yielded 8 CREs associated with F9, GP1BB, HBG1, IRF6, LDLR, MSMB, PKLR, and TERT. We evaluated each CRE using the same RNA tracks as in the Borzoi study: K562 for F9, GP1BB, HBG1, and PKLR; adrenal gland for LDLR, MSMB, and TERT; and keratinocyte, fibroblast, and epithelial tracks for IRF6.

For promoter variants, we additionally assigned each mutation to one of four categories: over-expression, under-expression, none, or unassigned, according to the classification defined by Jaganathan et al. [36].

For single-ISM, each variant within the CRE was evaluated individually as described above. For Multi-ISM, we performed Multi-ISM across the full 500 kb mutagenesis window centered on each gene and extracted the variant-effect estimates corresponding to each CRE.

### 5.9 GTEx v8 fine-mapped eQTL benchmark

We evaluated Multi-ISM and single-ISM on fine-mapped eQTLs from GTEx v8. We used SuSiE fine-mapping summary statistics and posterior probabilities generated in previous studies [1, 49]. Following the Borzoi evaluation protocol, we retained only single-nucleotide variants with posterior inclusion probability (PIP) *≥* 0.9 as high-confidence causal eQTLs across 26 GTEx tissue groups with matched Borzoi RNA-seq output tracks. Intersecting with the 5,000 genes for which Multi-ISM maps were computed yields 6,712 variant–gene–tissue triplets (1,778 genes, 2,971 unique variants) as the final benchmark set. We assess sign prediction accuracy and AUROC for classifying the direction of the fine-mapped eQTL effect (positive vs. negative posterior beta).

#### Single-ISM scoring

For each variant, we estimated its effect individually using the same procedure as single-ISM described above. Here, single-ISM refers to single variant effect estimation rather than true exhaustive mutagenesis. Effects were averaged across Borzoi RNA tracks matched to the corresponding GTEx tissue.

#### Multi-ISM scoring

For each variant, we extracted the corresponding variant effect estimate from the precomputed 500 kb Multi-ISM map for the target gene and averaged the estimates across Borzoi RNA tracks matched to the corresponding GTEx tissue. Variants with Multi-ISM coefficients of zero therefore received a predicted effect of exactly zero.

#### Borzoi and AlphaGenome detection-limit

For Supp. Fig. S1A, we also used fine-mapped GTEx v8 eQTLs with PIP *≥* 0.9. But we focused on the overlap of variant–gene–tissue triplets from the published Borzoi and AlphaGenome studies [1, 2]. For Borzoi, we used ensemble predictions across four replicate models. For AlphaGenome, we downloaded the published prediction scores from https://storage.googleapis.com/alphagenome/evals/eqtl_variant_borzoi_coefficient_human_predictions.feather, restricted to test set variants.

### 5.10 CRISPR FlowFISH Enhancer Prioritization Benchmark

We benchmarked enhancer prioritization using the same CRISPRi-FlowFISH dataset as previous work [29], which quantified expression changes for 53 genes in K562 cells after CRISPRi perturbation of candidate enhancers. Each method scored every enhancer–gene pair within the model input context window, and performance was evaluated using AUROC and AUPRC, stratified by enhancer–TSS distance: *<* 15 kb, 15–45 kb, 45–98 kb, and 98–250 kb.

#### Enformer

We used the precomputed scores from the Borzoi manuscript [1, 50].

#### Borzoi

We applied the same gradient-based scoring and normalization procedure described in the Borzoi manuscript to the Borzoi fold 0 model [1].

#### Flashzoi

We followed the same procedure as Borzoi [14]. We used the fold 0 model to compute gradients of the K562 RNA-seq prediction with respect to the input sequence. Specifically, for each gene, we computed gradients with respect to the sum of all output bins overlapping the gene’s exons and averaged across K562 RNA-seq tracks. We mean-centered the gradients across nucleotide channels and retained the gradient of the reference nucleotide. To score candidate enhancer regions, we applied Gaussian smoothing with *σ* = 300 bp to the absolute gradient values, then normalized the smoothed scores by mean across the entire region to account for differences between genes. The normalized score at the center base was used as the final score for each enhancer.

#### Multi-ISM

We ran Multi-ISM on each of the 53 genes. The resulting variant effect scores were averaged across 71 K562 RNA target tracks and converted into a Multi-ISM attribution vector by mean-centering across the four nucleotide channels and retaining the score of the reference nucleotide. To prioritize candidate enhancers, we used the Gaussian-smoothed (*σ* = 300 bp) absolute attribution value at the enhancer center. Cross-gene normalization was not applied because ISM scores are measured in units of log fold change and are therefore directly comparable across genes.

### 5.11 Multi-ISM nucleotide attribution score

For each gene, Multi-ISM predicts the log-fold-change effect of the three possible nucleotide substitutions at every genomic position and for each prediction track. For a given position and track, let *s*_alt1_, *s*_alt2_, and *s*_alt3_ denote the three substitution effects directly estimated by Multi-ISM. We define the contribution of the reference nucleotide as

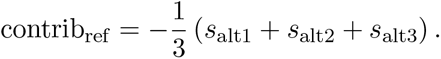

This quantity represents the negative mean log-fold-change induced by substituting the reference nucleotide with each of the three alternatives. It is, up to a constant scale factor, equivalent to the conventional reference-nucleotide attribution obtained after mean-centering across the four nucleotide channels.

Using these Multi-ISM derived nucleotide attribution scores, we constructed a position-by-track matrix *R* ∈ ℝ*^L×T^* for each gene, where *L* is the genomic window length and *T* is the number of prediction tracks.

### 5.12 Multi-ISM based characterization of regulatory complexity

We used effective rank to summarize gene regulatory complexity from the full Multi-ISM nucleotide-contribution matrix *R* ∈ ℝ*^L×T^*. To characterize regulatory variation along the track axis, we decomposed *R* using singular value decomposition:

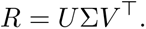

We then computed the effective rank of *R* using the standard entropy-based definition [51]. Given the nonzero singular values *σ*_1_*,…, σ_k_*, we normalized the singular-value spectrum as 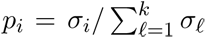 and calculated

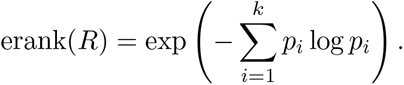

Effective rank summarizes how concentrated the singular-value spectrum of R is, and we use it as a relative measure of pattern diversity across tracks rather than as an absolute count of independent regulatory patterns. A low effective rank indicates that the attribution matrix is dominated by a small number of patterns shared across tracks, whereas a high effective rank indicates that multiple distinct track-level patterns contribute substantially.

To localize this complexity, we further computed the effective rank within subsets of positions defined by genomic annotation. Each position was assigned to exactly one category based on GENCODE annotations, using the following priority order: splice site (within 10 bp of an intron–exon boundary), coding (CDS), 3’-UTR, promoter (within 2 kb of any TSS), and distal non-coding (all remaining positions). We report the effective rank across all positions and separately within each annotation category.

We correlated effective rank with a few independent measurements of regulatory complexity.

#### GTEx tissue-specificity index

was quantified using the *τ* index calculated from GTEx v8 RNA-seq data [28]. For each gene, expression in tissue *i* was summarized across samples as *x_i_* = log_2_ (median(TPM*_i_*) + 1). For a gene measured across *N* tissues, *τ* was calculated as

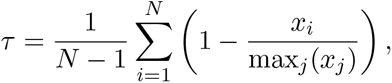

where max*_j_*(*x_j_*) denotes the maximum log-transformed median expression across tissues. The *τ* index ranges from 0, indicating uniform expression across tissues, to 1, indicating expression restricted to a single tissue.

#### rE2G enhancer linkage

was quantified using ENCODE-rE2G predicted enhancer–gene links, downloaded from the ENCODE portal as one thresholded prediction file per biosample [29]. We used five cell lines: K562 (ENCFF976OKL), GM12878 (ENCFF978BWG), HepG2 (ENCFF775WBY), A549 (ENCFF539BQT), and MCF-7 (ENCFF213ORL). For each cell line, we retained only distal enhancer links by excluding self-promoter elements (isSelfPromoter == False) and counted the number of remaining links assigned to each target gene. We correlated effective rank with the average number of rE2G links per gene across all five cell lines.

#### Promoter basal strength and responsiveness

were obtained from the 21E *×* 500P MPRA performed by Tan et al. [27]. After filtering, the dataset contained measurements for 269 distinct promoters in K562 cell lines. We computed Multi-ISM scores for 248 of the corresponding genes. *P*_basal_ was defined as the mean log_2_ expression of each promoter when paired with inactive control enhancers, whereas *P*_resp_ was defined as the zero-intercept slope relating enhancer-induced expression changes at that promoter to the average activity of each enhancer.

#### Candidate cis-regulatory element (cCRE) coverage

of each gene is defined as the amount of candidate regulatory sequence within a 500-kb window centered on the gene using the Version 3 ENCODE SCREEN Registry of cCREs (1,063,878 elements; ENCODE Project Consortium) [52]. To exclude promoter-proximal sequence, we removed regions within *±*1 kb of all annotated transcription start sites (TSSs) for the gene. `cre_bp` is defined as the total number of non-overlapping base pairs covered by the remaining distal cCREs within the 500-kb window.

**The gene constraint coefficient (***s*_het_) is downloaded from Zeng et al [53]. It is a genelevel estimate of the fitness cost of carrying a heterozygous loss-of-function variant, inferred from population genetic patterns of LoF variation under an explicit selection model.

#### Functional and disease gene sets

Functional gene sets were obtained from MSigDB v2025.1.Hs.

We used the C5 terms `GOBP_RIBOSOME_BIOGENESIS, GOBP_TRANSLATION, GOBP_DNA_REPAIR, GOBP_CHROMATIN_REMOD_GOBP_EMBRYO_DEVELOPMENT, GOBP_NEURON_MIGRATION, and GOBP_CELL_FATE_COMMITMENT`, together with the C2:CGP set `HOUNKPE_HOUSEKEEPING_GENES`. Disease gene sets were obtained from the MacArthur Lab resource (https://github.com/macarthur-lab/gene_lists) and included autosomal-dominant (709 genes), autosomal-recessive (1,183 genes) [54, 55], and haploinsufficient genes (294 genes) [56]. Gene-set assignments were non-exclusive, and each set was compared with the genome-wide background (“All genes”) using a two-sided Mann–Whitney *U* test followed by FDR correction.

### 5.13 Multi-ISM-based mutational sensitivity analysis

For each gene and annotation category (promoter, splice site, coding, 3*^′^* UTR, other noncoding, and all positions), we summarized the Multi-ISM effects using five statistics: p99 Multi-ISM, 99th percentile of absolute non-zero effects; mean Multi-ISM, mean absolute non-zero Multi-ISM effect; max Multi-ISM, maximum absolute effect; effective rank, effective rank of the position-by-track matrix; nonzero Multi-ISM, fraction of non-zero matrix entries.

#### Partial correlation analysis

We tested the association between each Multi-ISM statistic and gene constraint, log_10_(*s*_het_). Partial Spearman correlations were computed after adjusting for six gene-level covariates: log mean expression, tissue-specificity *τ*, exon count, CDS length, gene span, and promoter GC content. All variables were rank-transformed, the ranked Multi-ISM statistic and ranked constraint were separately regressed on the ranked covariates, and the Pearson correlation between the two residuals was reported as the partial Spearman correlation. Genes with missing values were excluded per annotation category.

#### Cross-validated prediction of gene constraint

To test whether Multi-ISM statistics improved prediction of log_10_(*s*_het_) beyond the same six gene-level covariates, we evaluated linear regression models using 10-fold cross-validation. Features were standardized using the training fold only and then applied to the held-out fold.

For each annotation category, we compared four feature sets: gene covariates alone; covariates plus log effective regulatory rank; covariates plus the 99th-percentile absolute effect; and covariates plus both Multi-ISM features. Performance was evaluated by Spearman correlation, and *R*^2^ using pooled out-of-fold predictions.

### 5.14 Personal Gene Expression Prediction

#### 5.14.1 Preparation of GTEx whole-blood expression and genotype data

The GTEx v8 Whole Blood normalized expression matrix and matching eQTL covariate files were downloaded from the GTEx Portal (https://gtexportal.org/home/downloads/adult-gtex/qtl). For each gene, we regressed the normalized expression on the covariates and z-scored the residuals per gene across individuals. This covariate-corrected expression profile is the final gene expression table we predict. Individual WGS genotypes were obtained from dbGaP under accession phs000424.v8. A total of 670 donors had both expression and genotype data.

#### 5.14.2 Whole blood expression outliers

To call expression outliers in whole blood, we followed the procedure of Ferraro et al [42]. For each gene, we regressed the covariate-corrected expression on its lead significant cis-eQTL variant and z-scored the residuals across individuals. The resulting per-gene z-score is our expression-outlier score for each individual, and we define an expression outlier as any (gene, individual) pair with |*z*| > 3

#### 5.14.3 Multi-ISM variant-burden scoring

For each gene of interest, Multi-ISM returns a mutational table giving the effect of every possible mutation in a 500-kb window around the gene center on all Borzoi RNA and CAGE tracks. We summarize a variant’s effect in whole blood as its mean ISM score across the GTEx whole-blood RNA targets. Because Multi-ISM is estimated under an additivity assumption, we score a haplotype by looking up its ALT alleles within the 500-kb window and summing their effects, yielding a predicted log fold-change relative to the reference haplotype; an individual’s burden score is the mean of the two haplotype scores. For each individual we computed a Multi-ISM rare-variant burden score on variants with MAF *<* 5% (MISM-rare), and additionally, a Multi-ISM variant burden score aggregating variants across the full allele-frequency spectrum (MISM-all).

#### 5.14.4 Elastic-net models for genotype-based expression prediction

For each gene, we built genotype-based prediction models of whole-blood expression using a PrediXcan-style elastic-net approach. The 670 donors were divided into a fixed 80% training set and 20% test set, with the same split used across all genes and all models. Models were fit with scikit-learn’s ElasticNet and ElasticNetCV.

##### EN-common (common-variant elastic net)

For each individual, the input features were the alt-allele dosages (0/1/2) of all common variants (MAF *≥* 0.05) within the same 500-kb window centered on the gene as used by Multi-ISM. For each gene, we fit an elastic-net model on the training set with five-fold cross-validation. We considered *ℓ*_1_ *∈ {*0.5, 0.6, 0.7, 0.8, 0.9, 1.0*}* and penalty parameters *α ∈ {*10*^−^*^5^, 10*^−^*^4^, 10*^−^*^3^, 10*^−^*^2^, 10*^−^*^1^, 1, 10, 100*}*. The (*α, ℓ*_1_-ratio) combination that minimized the cross-validated mean squared error was selected, and the model was then refit on the full training set. Model performance was evaluated in the held-out individuals using Pearson’s *R* between predicted and observed z-scored expression. This is the standard PrediXcan-style genotype-based expression prediction setting used as our baseline, which we refer to as the *common-variant elastic net*.

##### EN-common + MISM (Multi-ISM-augmented elastic net)

In addition to the genotype features used in EN-common, we added a z-score-normalized Multi-ISM burden computed by aggregating the predicted effects of variants with MAF *<* 0.05; we refer to this feature as MISM. The elastic-net model was then refit with MISM included as an additional feature. For genes with a non-constant EN-common prediction, we reused the EN-common hyperparameters (*ℓ*_1_ ratio and *α*) to isolate the contribution of the added MISM feature. For genes with a constant EN-common prediction, for which all coefficients were shrunk to zero and Pearson’s *R* was undefined, we reran `ElasticNetCV` over the original hyperparameter grid because the inherited *α* would otherwise also shrink the MISM coefficient to zero. We refer to this model as the *Multi-ISM–augmented elastic net*.

##### EN-common + permuted MISM

This model uses the same design and training procedure as EN-common + MISM, except that the MISM feature is randomly permuted across individuals before being appended as the extra feature. This permutation null serves as a baseline to isolate how much of any predictive gain from MISM reflects genuine signal rather than the added model capacity of one extra feature.

##### EN-common + MISM-all

This model uses the same design and training setup as EN-common + MISM, except that the MISM feature is replaced by MISM-all, a Multi-ISM burden aggregated over variants across the full allele-frequency spectrum. Comparing MISM-all with MISM tests whether including common variants in the Multi-ISM burden provides additional benefit beyond the burden from variants with MAF *<* 0.05.

##### EN-0.01

This model follows the same design as EN-common, except that the genotype feature set is expanded to include variants with MAF *≥* 0.01 within the 500-kb window. For each gene, hyperparameters were selected by five-fold cross-validation, and the model was then refit on the full training set.

##### EN-0.01 + MISM

We added the z-score normalized MISM burden as an additional feature to the EN-0.01 genotype design matrix, following the same training setup (reuse EN-0.01 hyperparameters for defined baselines, and re-run ElasticNetCV for undefined baselines).

##### EN-all

This model follows the same design as EN-common, except that the genotype feature set is expanded to all variants (no MAF threshold) within the 500-kb window. For each gene, we selected hyperparameters by five-fold cross-validation and then re-fit on the full training set.

##### EN-all + MISM

We added the z-score normalized MISM burden as an additional feature to the EN-all genotype design matrix, following the same training setup (re-use EN-all hyperparameters for defined baselines, and re-run ElasticNetCV for undefined baselines).

## 6 Data and code availability

The Multi-ISM mutational maps generated in this study comprise nucleotide-resolution variant-effect predictions across 500-kb gene-centered windows for 5,000 protein-coding genes. The maps are available through an interactive web server at https://multi.ism.research.calicolabs.com/ and as downloadable files from the GCS bucket gs://calico-public-multi-ism-bucket/. GTEx v8 normalized expression and eQTL summary data are publicly available through the GTEx Portal (https://gtexportal.org/). Fine-mapped eQTLs were obtained from Wang et al. [49]. Individual-level GTEx whole-genome sequencing genotypes were obtained through dbGaP under accession phs000424.v8 and remain subject to controlled-access requirements. The CAGI5 saturation-mutagenesis data were obtained from Kircher et al. [48]. The CRISPRi-FlowFISH enhancer–gene benchmark was obtained from Gschwind et al. [29]. ENCODE candidate cis-regulatory elements were obtained from the ENCODE SCREEN Registry [52], rE2G enhancer–gene links from Gschwind et al. [29], and promoter responsiveness measurements from Tan et al. [27]. The Mendelian disease-gene set was derived from the OMIM-based curation distributed by Minikel et al. [47].

Multi-ISM scripts are available at https://github.com/calico/multi-ism.

## 8 Acknowledgments

This work was funded by Calico Life Sciences LLC. The funder had no role in study design, data collection, or analysis. Publication of the manuscript was approved after an internal scientific review process. We thank Joe Arron, David Wang, Qingbo Wang, Jason Tan, and Anya Korsakova for helpful discussions and valuable feedback.

---

**Figure S1:**
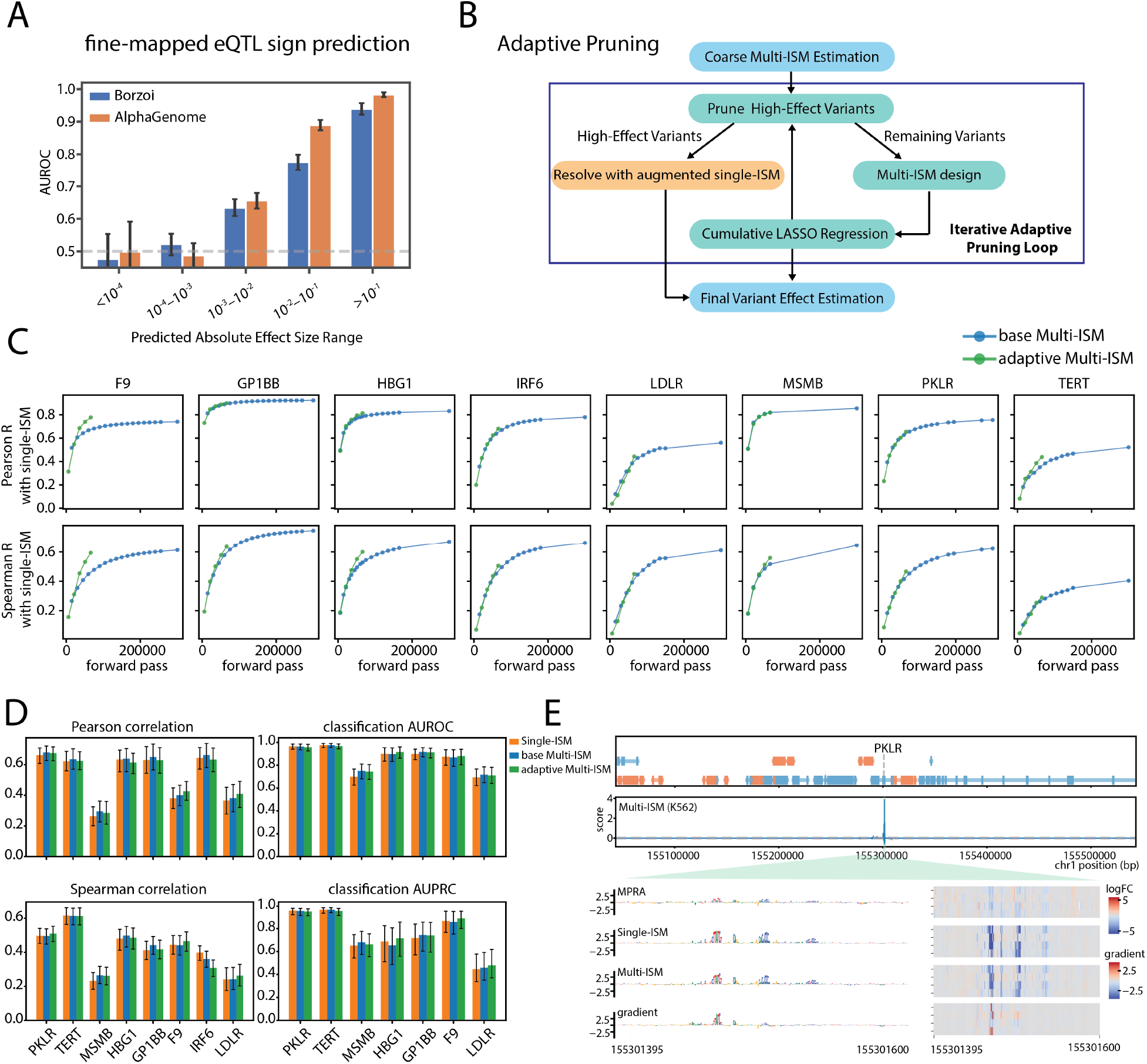
Multi-ISM performance on the CAGI5 MPRA benchmark. **A,** Sign prediction AUROC of Borzoi and AlphaGenome for fine-mapped GTEx v8 eQTLs (PIP *≥* 0.9, n=5,593), stratified by predicted variant effect magnitude. Predictive performance approaches random for variants with very small predicted effects. Error bars indicate 95% confidence intervals estimated from 1,000 bootstrap resamples. **B,** Schematic of adaptive Multi-ISM. After an initial coarse Multi-ISM estimate, high-effect variants are identified, re-estimated individually across four randomly mutated sequence backgrounds, and removed from subsequent mutation designs and regression fits. Additional Multi-ISM measurements are then used to refine the remaining unresolved variants, and the procedure is repeated iteratively. More details see **Methods**. **C,** Agreement of base and adaptive Multi-ISM with single-ISM across the eight CAGI5 MPRA loci as a function of computational budget, measured by Pearson’s and Spearman’s correlation. The comparison includes variants within the central 10-kb window of each gene with absolute single-ISM effects greater than 0.01. **D,** Performance on the CAGI5 MPRA benchmark for single-ISM, base Multi-ISM, and adaptive Multi-ISM. Pearson and Spearman correlations quantify agreement with experimentally measured variant effects; AUROC and AUPRC assess classification of promoter variants using the over-expression and under-expression labels from Jaganathan et al. [36]; the IRF6 enhancer element is unlabeled. Error bars indicate 95% confidence intervals estimated by bootstrap resampling. **E,** Top, adaptive Multi-ISM attribution scores across the 500 kb region surrounding PKLR in K562. In gene tracks, orange and blue denote plus- and minus-strand genes. Bottom, CAGI5 MPRA measurements, single-ISM, adaptive Multi-ISM and gradient attribution scores near PKLR promoter. Nucleotide-level substitution effects are shown at right for the indicated region; for gradient, nucleotide-level gradient scores are shown.

**Figure S2:**
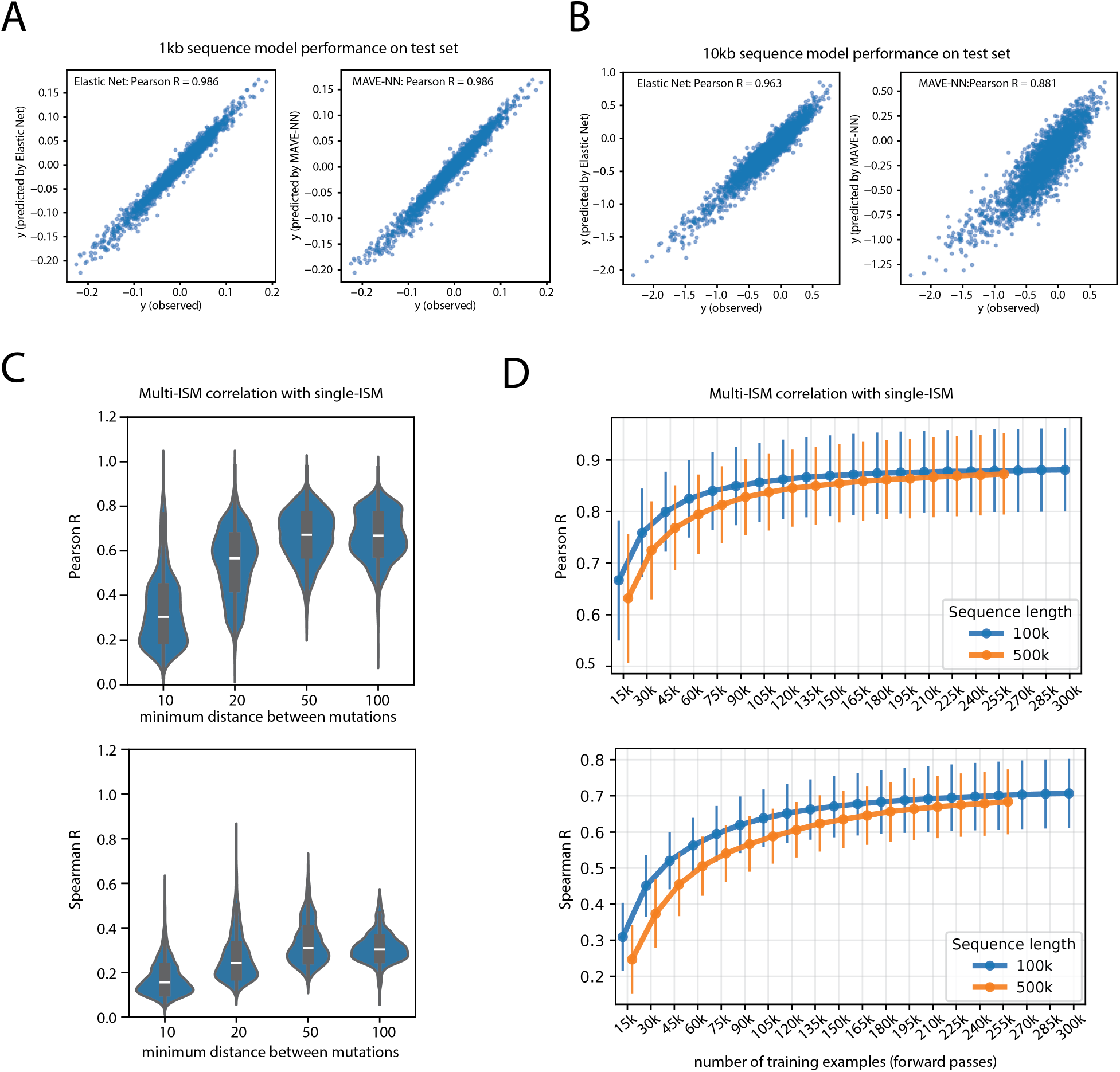
Sensitivity of base Multi-ISM design. **A,** Comparison of linear regression and MAVE-NN for modeling base Multi-ISM measurements over a 1kb mutagenesis region at the *GATA1* locus with a minimum distance of 50bp between mutations (**Methods**). Predicted versus observed GATA1 log-foldchange are shown for held-out multi-mutant sequences using elastic-net regression with Celer (left) or MAVE-NN (right). Both models achieve Pearson’s *R* = 0.986. **B,** Same comparison as in **A** over a 10 kb mutagenesis region. Elastic-net regression with *ℓ*_1_ ratio = 1 (LASSO) achieves Pearson’s *R* = 0.963, compared with *R* = 0.881 for MAVE-NN. **C,** Effect of minimum mutation spacing on base Multi-ISM performance at the *GATA1* locus. Performance evaluated by correlation with single-ISM over the central 10 kb of the window (**Methods**). Violin plots show the distribution across model output tracks of Pearson (top) and Spearman (bottom) correlations with single-ISM, using a fixed budget of 15,000 forward passes. Performance is reduced at 10–20 bp spacing, whereas 50 and 100 bp perform comparably well. **D,** Base Multi-ISM performance as a function of computational budget and sequence length. Points show the median Pearson (top) and Spearman (bottom) correlation with single-ISM across output tracks for 100 kb and 500 kb windows; error bars indicate *±*1 standard deviation. Correlation increases with additional measurements, with the 100 kb window performing better at low budgets and the two window sizes converging at higher budgets.

**Figure S3:**
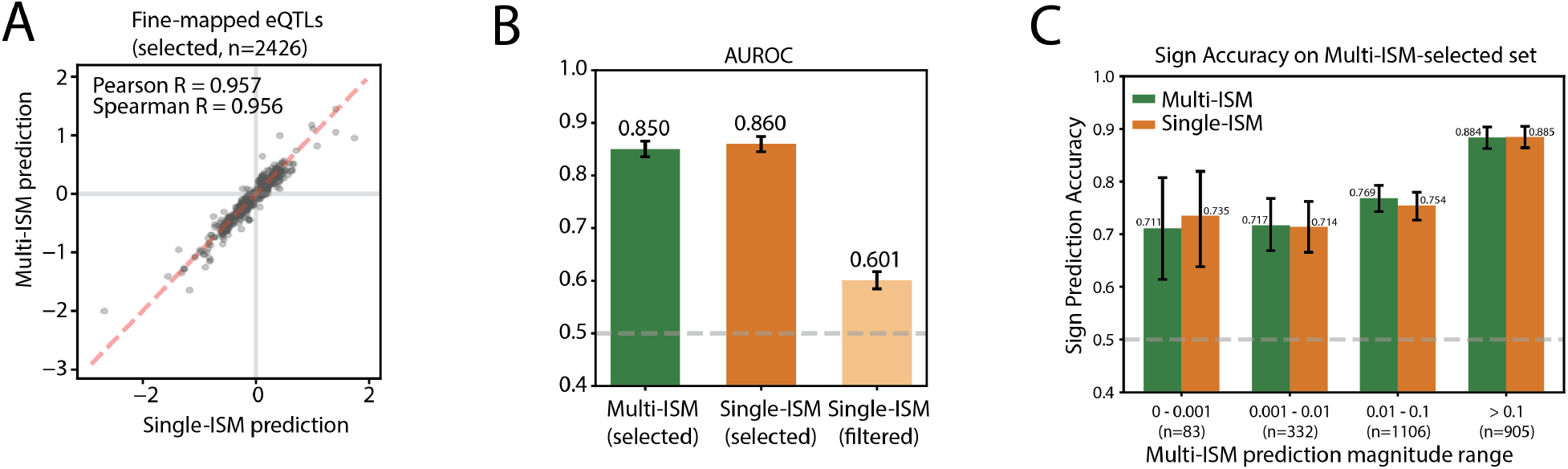
Multi-ISM variant effect prediction performance on GTEx eQTL. **A,** Correlation between Multi-ISM and single-ISM predictions for the 2,426 fine-mapped GTEx v8 eQTL variant–gene–tissue triplets (PIP *≥* 0.9) assigned non-zero effects by Multi-ISM (Pearson’s *R* = 0.957; Spearman’s *R* = 0.956). **B,** AUROC for predicting the direction of fine-mapped eQTL effects. Multi-ISM and single-ISM show similar performance on variants selected by Multi-ISM, whereas variants filtered by Multi-ISM have substantially lower predictive performance when scored by single-ISM. Error bars indicate 95% confidence intervals estimated from 1,000 bootstrap resamples. **C,** Sign-prediction accuracy for Multi-ISM-selected variants, stratified by Multi-ISM predicted effect magnitude and scored using either Multi-ISM or single-ISM. Numbers above bars indicate accuracy, and numbers below each bin indicate the number of variants evaluated.

**Figure S4:**
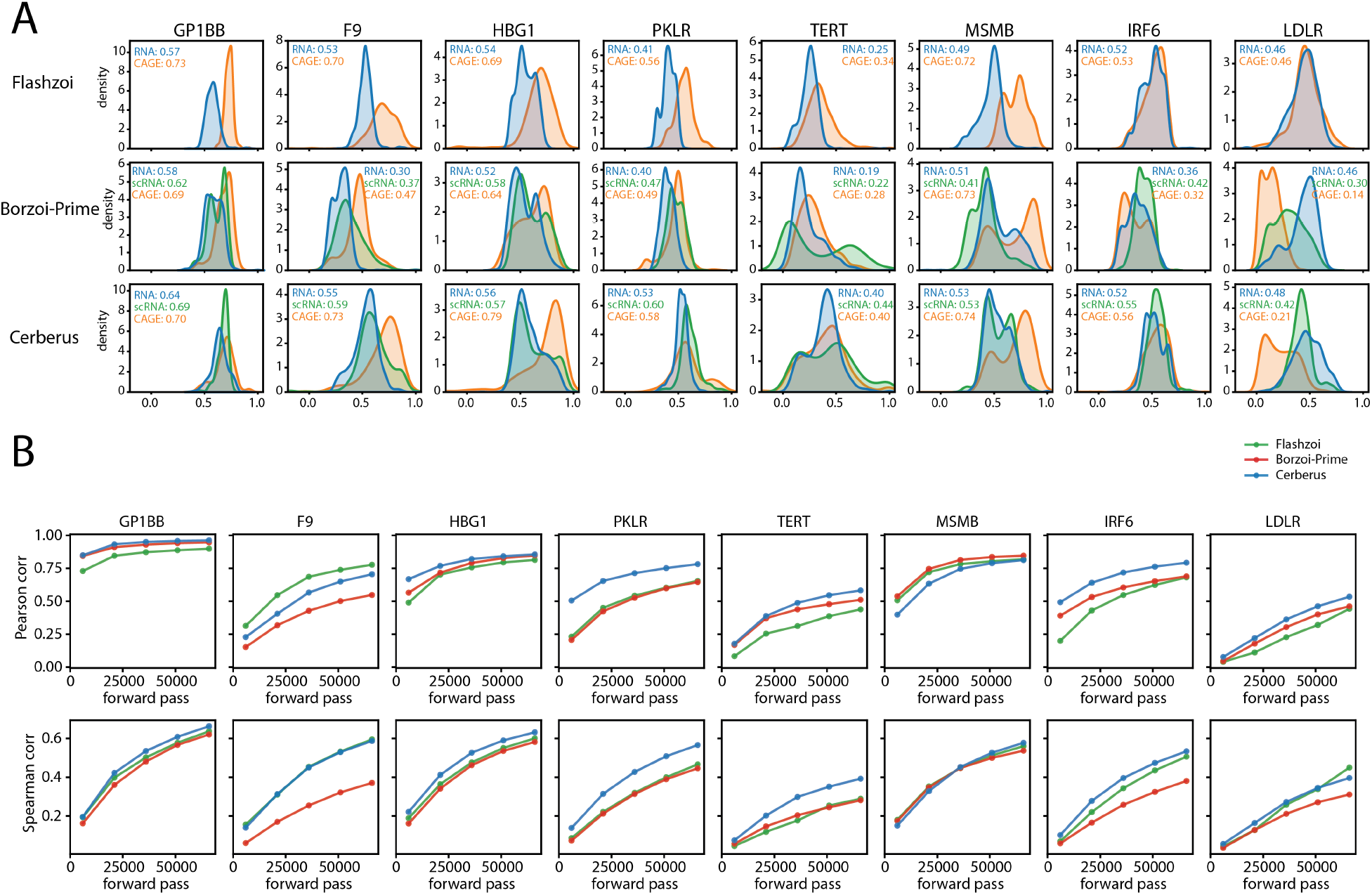
Multi-ISM recovers single-ISM effects across model architectures. **A,** Distribution of Spearman correlations between Multi-ISM and single-ISM variant-effect predictions across output tracks for eight example genes. Multi-ISM and single-ISM were evaluated using the same underlying model: Flashzoi (top), Borzoi-Prime (middle), or Cerberus (bottom). The comparison includes variants within the central 10-kb window with absolute single-ISM effects greater than 0.01. Colors indicate CAGE, RNA, and scRNA output tracks. **B,** Agreement between Multi-ISM and single-ISM as a function of forward-pass budget for the same eight genes. Pearson (top) and Spearman (bottom) correlations are averaged across output tracks. The comparison includes variants within the central 10-kb window with absolute single-ISM effects greater than 0.01. Green, Flashzoi; red, Borzoi-Prime; blue, Cerberus.

**Figure S5:**
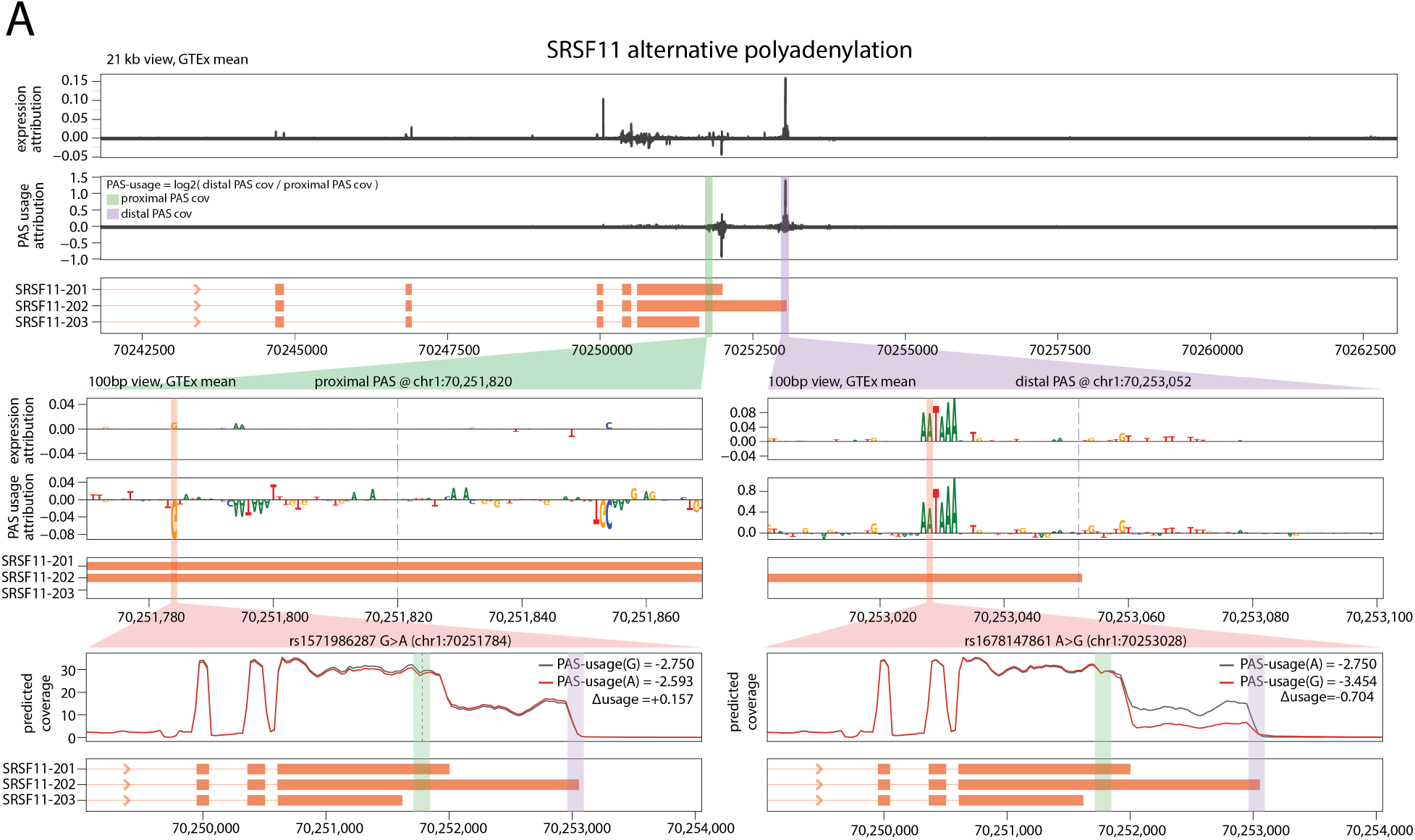
Multi-ISM PAS usage readout captures PAS-specific regulatory effects. **A,** The figure shows Multi-ISM attribution for expression (standard readout, defined as the log of summed exonic coverage) and for alternative polyadenylation site usage readout (PAS usage, defined as the log ratio of predicted coverage at the distal PAS to coverage at the proximal PAS). The 21 kb view shows attribution across the SRSF11 locus, with the proximal PAS at chr1:70,251,820 and the distal PAS at chr1:70,253,052 highlighted. The 100 bp views zoom in on sequence features surrounding each PAS, and the bottom tracks show Borzoi-predicted coverage for the two alleles of the highlighted variants. Left, rs1571986287 G*>*A lies upstream of the proximal PAS within a UGU/UGUA element with noticeable PAS-usage attribution but little expression attribution. The negative PAS-usage attribution indicates that mutations at this position are predicted to increase the distal-to-proximal usage ratio, consistent with reduced usage of the proximal PAS. In the predicted coverage tracks, the alt allele shows lower coverage before the proximal PAS and increased readthrough beyond it, indicating less efficient cleavage and polyadenylation at the proximal site. This increases the distal-to-proximal PAS-usage readout (Δusage = +0.157) while leaving overall transcript abundance largely unchanged. Right, rs1678147861 A*>*G disrupts the AAUAAA hexamer of the distal PAS and shows strong attribution for both PAS usage and expression. In the predicted coverage tracks, the alt allele produces a marked reduction in coverage leading up to the distal PAS, consistent with reduced abundance of the distal isoform. This decreases the distal-to-proximal PAS-usage readout (Δusage = *−*0.704) and is accompanied by reduced predicted expression.

**Figure S6:**
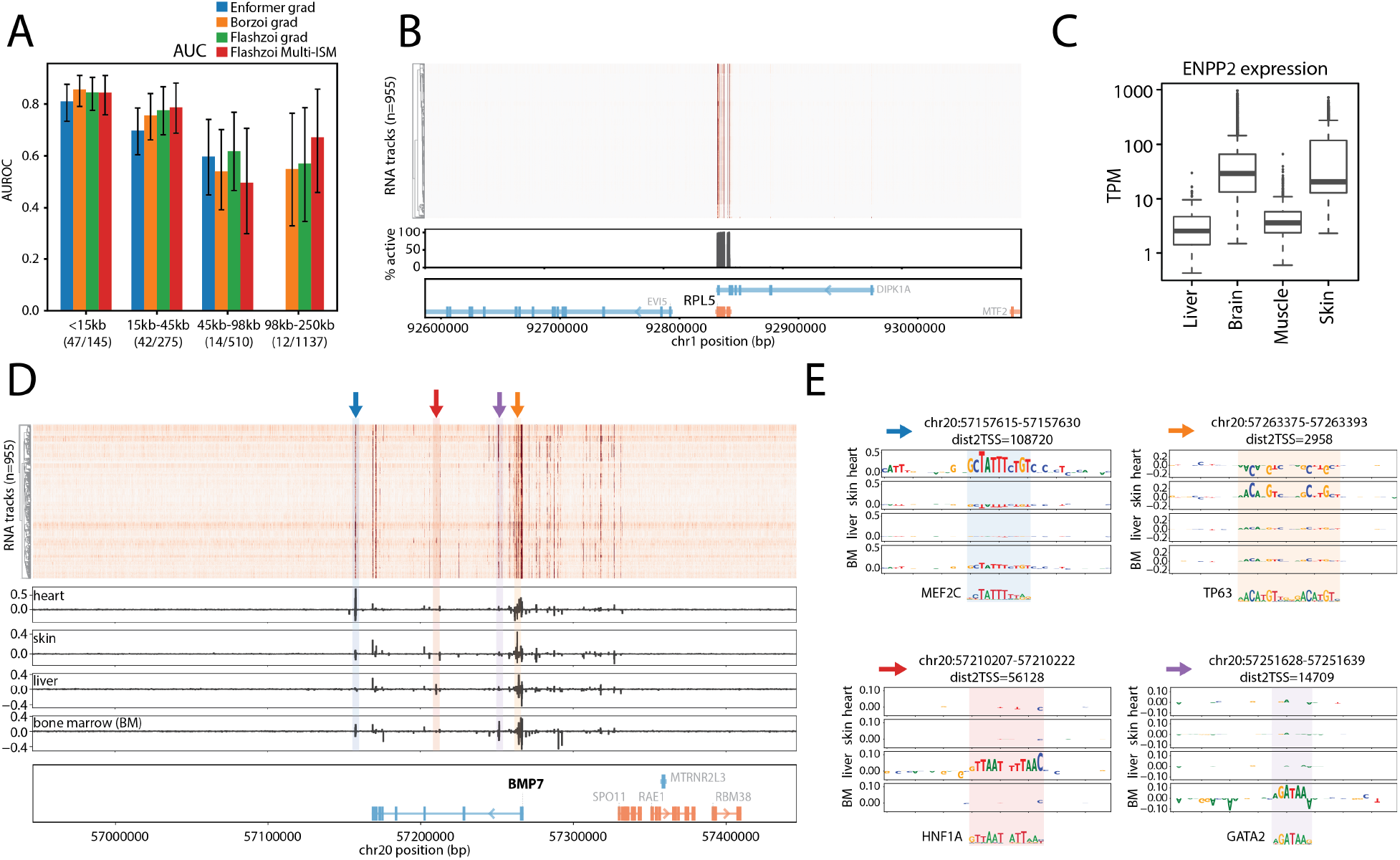
Additional evaluation of tissue-specific regulatory architecture identified by Multi-ISM. **A,** AUROC on the K562 CRISPRi-FlowFISH enhancer prioritization benchmark across 53 genes, stratified by enhancer–TSS distance. Enformer scores are results reported in the Borzoi study; Borzoi gradients, Flashzoi gradients, and Multi-ISM use fold 0 for matched comparison. Numbers below each bin indicate positive/total enhancer–gene pairs tested. Error bars indicate 95% confidence intervals estimated from 1,000 bootstrap resamples. **B,** Multi-ISM regulatory map across the 500-kb window surrounding *RPL5*. Top, track-by-position attribution heatmap across 955 RNA tracks, with each 100-bp bin summarized by its maximum absolute attribution. Tracks are ordered by hierarchical clustering (dendrogram at left). Bottom, percentage of tracks in which each 100-bp bin is active, defined as a maximum absolute attribution *>* 0.05. *RPL5* shows a compact regulatory architecture dominated by promoter- and gene-proximal sequence. **C,** GTEx expression of *ENPP2* in liver, brain, muscle, and skin, the four tissues examined in Fig. 4B–D. *ENPP2* is expressed in all four tissues, although expression levels vary. Boxes show the median and interquartile range. **D,** Multi-ISM regulatory map across the 500-kb window surrounding *BMP7*. Top, track-by-position attribution heatmap across 955 RNA tracks, with each 100-bp bin summarized by its maximum absolute attribution. Tracks are ordered by hierarchical clustering (dendrogram at left). Bottom, bp resolution tissue-specific Multi-ISM attribution profiles for heart, skin, liver, and bone marrow. Colored arrows and shaded regions mark four candidate tissue-selective regulatory elements examined in (E). In gene tracks, orange and blue denote plus- and minus-strand genes. **E,** Base-pair-resolution Multi-ISM attribution at the four candidate *BMP7* regulatory elements highlighted in (D), shown in representative tissue tracks together with matched transcription-factor motifs.

**Figure S7:**
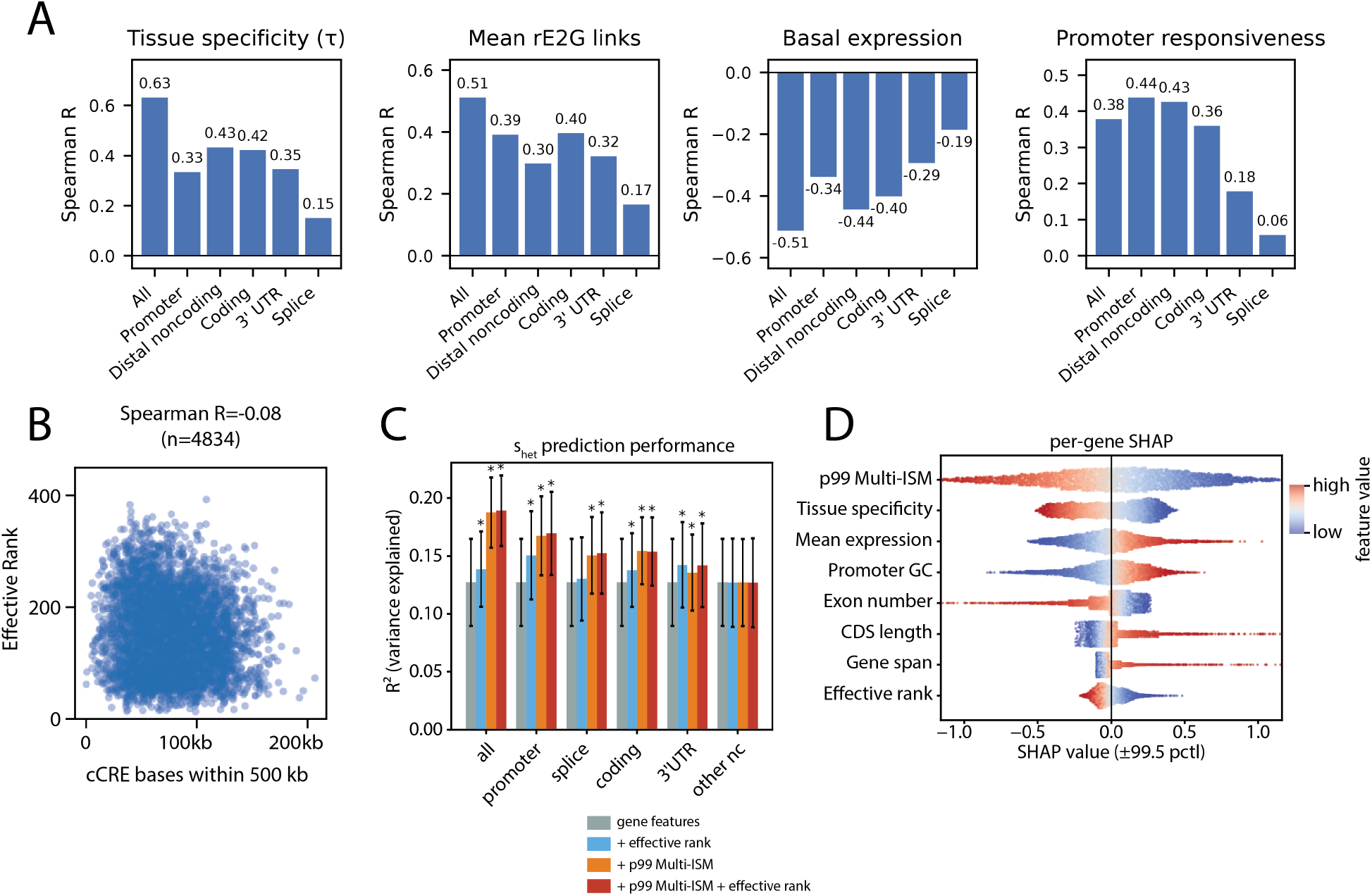
Regulatory complexity and mutational effect magnitude show distinct relationships with gene constraint. **A,** Spearman correlations between Multi-ISM effective rank metric and GTEx tissue specificity (*τ*), number of ENCODE rE2G links (averaged over 5 cell lines), basal promoter expression, and promoter responsiveness from K562 MPRA. Effective rank is computed using the full 500-kb attribution matrix (All) or separately within promoter, distal non-coding, coding, 3*^′^*-UTR, and splice-site regions. Genomic categories are defined from GENCODE annotations (**Methods**). **B,** Effective rank versus the total number of base pairs overlapping ENCODE candidate cis-regulatory elements (cCREs) within the 500-kb gene-centered window. Spearman correlation and sample size are shown. **C,** Cross-validated prediction of *s*_het_ using gene features alone (gray), plus effective rank (blue), plus p99 mism (orange), or plus both (red), evaluated separately by annotation category. Bars show *R*^2^ between pooled held-out predictions and observed *s*_het_; error bars show standard deviation across folds. Asterisks indicate improvement over gene features alone by one-sided Wilcoxon signed-rank test across folds (*P <* 0.05). **D,** SHAP values for held-out genes from the cross-validated full *s*_het_ prediction model (gene features + effective rank + p99 mism). Each point represents a gene and is colored by its corresponding feature value (red, high; blue, low). SHAP values are clipped at the 0.5th and 99.5th percentiles for visualization.

**Figure S8:**
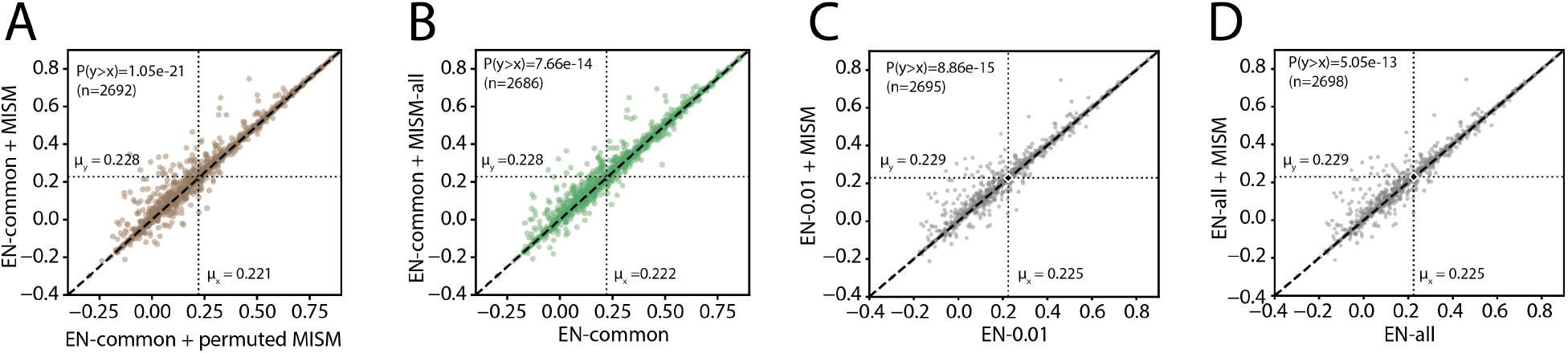
Additional baselines on Multi-ISM-augmented personalized expression prediction. Each point represents a gene, with performance measured by Pearson correlation between predicted and observed expression across held-out individuals. Dashed lines indicate *y* = *x*, and dotted lines indicate the mean performance of each model. *P* values are from one-sided Wilcoxon signed-rank tests. **A,** EN-common + MISM (*Multi-ISM–augmented elastic net*) compared with a control model in which the MISM burden was permuted across individuals. Genes with defined correlations in both models (2692) are shown. **B,** EN-common + MISM-all, which adds the Multi-ISM burden aggregated across variants of all allele frequencies as an additional feature to elastic net, compared with EN-common (*common variant elastic net*). Genes with defined correlations in both models (2686) are shown. **C,** EN-0.01 + MISM compared with EN-0.01, whose baseline elastic-net models use genotype features from variants with MAF *≥* 1%. Note the MISM burden is the same feature used in the Multi-ISM–augmented elastic-net model. Genes with defined correlations in both models (2695) are shown. **D,** EN-all + MISM compared with EN-all, whose baseline elastic-net models use genotype features from all variants regardless of allele frequency. Genes with defined correlations in both models (2698) are shown.

**Figure S9:**
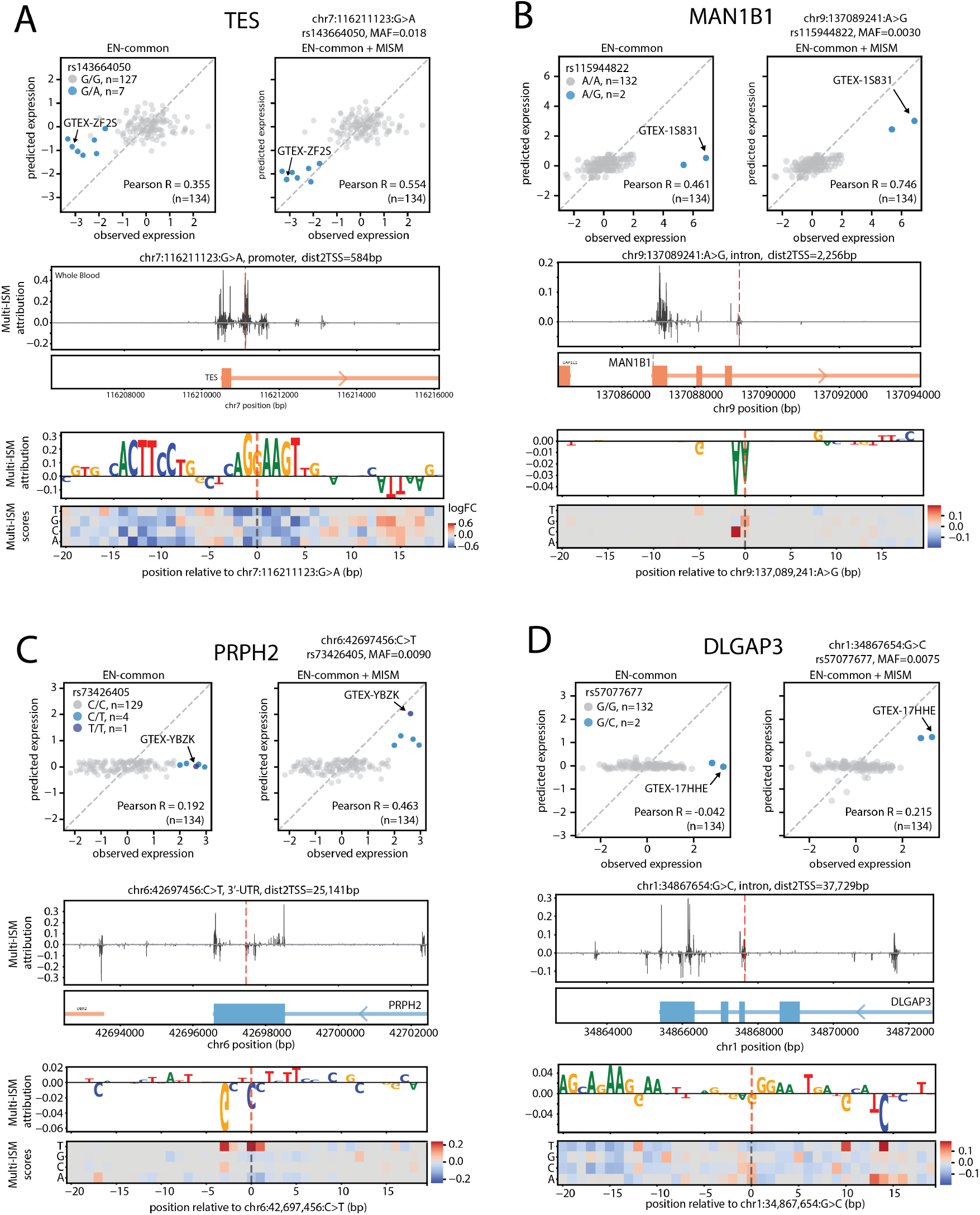
Additional examples of variants prioritized by Multi-ISM for personalized expression prediction. Each panel shows, from top to bottom, observed versus predicted whole-blood expression in held-out individuals using *common-variant elastic net* (EN-common) and *Multi-ISM–augmented elastic net* (EN-common+MISM), 10 kb local Multi-ISM attribution profile around the prioritized variant, and 40bp nucleotide-resolution Multi-ISM map shown as a seqLogo and heatmap. Variant carriers are highlighted in blue. **A,** *TES* promoter variant rs143664050 G*>*A (MAF = 0.018), associated with reduced expression in carriers; adding the Multi-ISM burden improves held-out Pearson correlation from *R* = 0.355 to *R* = 0.554. **B,** *MAN1B1* intronic variant rs115944822 A*>*G (MAF = 0.0030), annotated in ClinVar as creating a cryptic splice acceptor, associated with increased expression in carriers; Pearson correlation improves from *R* = 0.461 to *R* = 0.746. **C,** *PRPH2* 3*^′^*-UTR variant rs73426405 C*>*T (MAF = 0.0090), associated with increased expression in carriers; Pearson correlation improves from *R* = 0.192 to *R* = 0.463. **D,** *DLGAP3* intronic variant rs57077677 G*>*C (MAF = 0.0075), associated with increased expression in carriers; Pearson correlation improves from *R* = *−*0.042 to *R* = 0.215.

